# Nascent chromatin occupancy profiling reveals locus and factor specific chromatin maturation dynamics behind the DNA replication fork

**DOI:** 10.1101/398438

**Authors:** Mónica P. Gutiérrez, Heather K. MacAlpine, David M. MacAlpine

## Abstract

Proper regulation and maintenance of the epigenome is necessary to preserve genome function. However, in every cell division, the epigenetic state is disassembled and then re-assembled in the wake of the DNA replication fork. Chromatin restoration on nascent DNA is a complex and regulated process that includes nucleosome assembly and remodeling, deposition of histone variants, and the re-establishment of transcription factor binding. To study the genome-wide dynamics of chromatin restoration behind the DNA replication fork, we developed Nascent Chromatin Occupancy Profiles (NCOPs) to comprehensively profile nascent and mature chromatin at nucleotide resolution. While nascent chromatin is inherently less organized than mature chromatin, we identified locus specific differences in the kinetics of chromatin maturation that were predicted by the epigenetic landscape, including the histone variant H2A.Z which marked loci with rapid maturation kinetics. The chromatin maturation at origins of DNA replication was dependent on whether the origin underwent initiation or was passively replicated from distal-originating replication forks suggesting distinct chromatin assembly mechanisms between activated and disassembled pre-replicative complexes. Finally, we identified sites that were only occupied transiently by DNA-binding factors following passage of the replication fork which may provide a mechanism for perturbations of the DNA replication program to shape the regulatory landscape of the genome.

## Introduction

Chromatin organization is essential to maintain and regulate almost all aspects of genome function. The distribution and phasing of histone octamers on the DNA as well as the location of DNA binding proteins such as transcription factors define the regulatory landscape of the genome and govern transcription (Jiang and Pugh 2009; ENCODE Project Consortium 2012). The chromatin landscape is dynamic and responds to both developmental and environmental cues to modulate cell type specific gene expression programs. In addition to regulating gene expression, the local chromatin environment is also critical for other DNA templated processes such as DNA replication and repair (MacAlpine and Almouzni 2013; Dabin, Fortuny, and Polo 2016; Gutiérrez and MacAlpine 2016). Despite the central role of chromatin in genome function, every cell cycle the chromatin landscape must be disassembled ahead of the replication fork and then re-assembled behind the fork to preserve epigenetic memory.

Elegant genetic and biochemical experiments have elucidated many of the factors and mechanisms involved in the assembly of chromatin behind the DNA replication fork (Smith and Stillman 1989; Chang et al. 1997; Q. Li et al. 2008; Tyler et al. 1999; Schlesinger and Formosa 2000; Luk et al. 2007). The conserved chromatin assembly factor 1 (CAF-1), an H3-H4 histone chaperone, was first identified promoting histone deposition onto replicating SV40 DNA *in vitro* (Smith and Stillman 1989). CAF-1 is coupled to the replisome via an interaction with PCNA to ensure rapid histone deposition at the replication fork (Shibahara and Stillman 1999). Another H3-H4 histone chaperone, ASF1, cooperates with MCM2 at the fork to capture parental H3-H4 dimers which are then assembled into H3-H4 tetramers by CAF-1 for deposition on the nascent DNA (Huang et al. 2015; Richet et al. 2015; Sauer et al. 2017). The H2A/H2B histone chaperone, NAP-1, completes the assembly of the histone octamer on the DNA to form the nucleosome (Chang et al. 1997; Mosammaparast, Ewart, and Pemberton 2002). The ordered deposition of histone octamers behind the replication fork is critical for viability, genome stability and the maintenance of epigenetic state (Exner et al. 2006; Jasencakova et al. 2010; Cheloufi et al. 2015; Ishiuchi et al. 2015).

The assembly of nascent chromatin is tightly coupled to the replication fork. Early electron microscopy studies found a similar density of nucleosomes (“beads on a string”) on both the parental and nascent DNA strands (McKnight and Miller 1977), indicating that histone deposition and nucleosome formation must occur rapidly behind the replication fork. Consistent with these observations, reconstitution of replication-coupled assembly revealed that nucleosome assembly occurred within ~250 bp of the replication fork (Sogo et al. 1986; Cusick, Wassarman, and Depamphilis 1989; Gasser, Koller, and Sogo 1996). Despite the rapid deposition of the histone octamer behind the fork, nascent chromatin is differentially sensitive to nuclease digestion as compared to mature chromatin (>20 minutes post replication) (DePamphilis and Wassarman 1980; Klempnauer et al. 1980; Annunziato and Seale 1982; Stillman 1986), suggesting that nucleosomes are non-uniformly spaced in nascent chromatin. Together, these results underscore the complex and dynamic process by which chromatin matures following the rapid deposition of the histone octamer.

The organization of mature chromatin is dictated by many factors, including primary DNA sequence (Segal et al. 2006; Mavrich et al. 2008), the presence of pioneer factors (Bai et al. 2010; M. Li et al. 2015; Yan, Chen, and Bai 2018), and active transcription (Weiner et al. 2010). Conserved and stereotypical patterns of chromatin organization have emerged from micrococcal nuclease (MNase)-based studies of nucleosome positioning in a wide variety of eukaryotic organisms (Cui and Zhao 2012). Genes typically have well phased nucleosomes starting with the +1 nucleosome at the transcription start site (TSS) and proceeding into the gene body. Promoter regions are commonly marked by a nucleosome-free region (NFR) which are thought to accommodate regulatory factors. Similarly, well-positioned nucleosomes are observed flanking origins of DNA replication and their positioning is a determinant of origin function (Berbenetz, Nislow, and Brown 2010; Eaton et al. 2010; Belsky et al. 2015).

The study of nascent chromatin has been facilitated by the use of nucleoside analogs that allow for the affinity capture and purification of newly synthesized DNA. The enrichment of labeled nascent chromatin has been used in proteomic studies to identify proteins and protein networks associated with normal, stalled and collapsed replication forks (Sirbu et al. 2013). Similarly, others have described the maturation of post-translational histone modifications and the identification of new replisome factors important for maintaining genome stability (Alabert et al. 2014). While these proteomic studies provided a wealth of data on the network of proteins that ensure the stability and progression of the DNA replication fork and the temporal order in which chromatin modifications occur, they fail to reveal information about locus-specific differences in chromatin maturation.

Recent work by multiple groups have combined the power of 5-ethynyl-2-deoxyuridine (EdU) labeling of nascent DNA with MNase nucleosome mapping to ascertain the positioning of nucleosomes genome-wide in nascent and mature chromatin (Fennessy and Owen-Hughes 2016; Vasseur et al. 2016). Specifically, nascent DNA is labeled by a short pulse of EdU followed by a longer chase period to allow chromatin maturation. The EdU-labeled DNA is purified and subjected to next-generation sequencing following digestion with MNase. These studies focused on the chromatin maturation dynamics of nucleosomes within gene bodies and their rapid re-acquirement of nucleosomal organization and phasing, highlighting the role of transcription and histone chaperones in shaping the chromatin landscape. Importantly, studies in *Drosophila* found that the re-establishment of chromatin architecture at gene regulatory elements (eg. promoters and enhancers) was dependent on the re-association of transcription factors (Ramachandran and Henikoff 2016).

We have combined MNase epigenome mapping (Henikoff et al. 2011; Belsky et al. 2015) with EdU labeling of recently replicated DNA to generate nascent chromatin occupancy profiles (NCOPs) allowing us to holistically explore chromatin maturation dynamics in *S. cerevisiae*. We are able to resolve, at near nucleotide resolution, the maturation of nucleosomes and smaller DNA binding factors providing a factor agnostic view of chromatin assembly dynamics throughout the genome. We found that nascent chromatin was less organized than mature chromatin; however, there were locus specific differences in the maturation kinetics that were predicted by the epigenetic landscape. For example, poorly transcribed genes marked with the histone variant H2A.Z exhibited rapid chromatin maturation. Our results also confirm the role that site-specific DNA binding factors have in establishing chromatin organization and nucleosome positioning following passage of replication fork (Yadav and Whitehouse 2016; Yan, Chen, and Bai 2018). Strikingly, we also identified origin specific differences in chromatin maturation that were dependent on whether an origin initiated DNA replication or whether it was passively replicated from a neighboring fork, which may suggest an active mechanism to re-establish origin chromatin architecture at efficient origins. Finally, our factor agnostic approach to studying chromatin maturation revealed sites of transient occupancy by DNA-binding factors behind the replication fork, underscoring the plasticity of the chromatin landscape.

## Results

### Profiling nascent chromatin occupancy

We developed nascent chromatin occupancy profiles (NCOPs) to provide a factor-agnostic view of protein-DNA occupancy on newly synthesized DNA at nucleotide resolution. Specifically, we combined the power of labeling nascent DNA with the nucleoside analog EdU (Sirbu et al. 2011) with MNase-based epigenome mapping (Henikoff et al. 2011; Belsky et al. 2015; Ramachandran and Henikoff 2016). To demonstrate the sensitivity and specificity of NCOPs, we took advantage of the intra-S-phase checkpoint to specifically label newly synthesized DNA proximal to early origins of DNA replication. Briefly, yeast cells engineered to incorporate EdU (Viggiani and Aparicio 2006) were arrested in G1 by addition of the yeast mating hormone, α-factor, and subsequently released into media containing 200 mM hydroxyurea (HU) and 130 µM EdU. HU treatment depletes nucleotide pools which results in replication fork stalling and activation of the intra-S-phase checkpoint to prevent further origin activation (Santocanale and Diffley 1998; Shirahige et al. 1998). Only those sequences proximal (~10 kb) to early activating efficient origins will incorporate EdU into the nascent daughter strands. Following EdU labeling, chromatin was isolated and digested by MNase. Then, the EdU labeled DNA was biotin labeled by click chemistry prior to streptavidin affinity capture. Streptavidin-bound DNA was recovered and subjected to next-generation paired-end sequencing on the Illumina platform (Henikoff et al. 2011; Belsky et al. 2015)

We first generated chromosome-wide coverage plots of the sequencing depth to verify that EdU incorporation was specific and restricted to sequences proximal to early activating origins of DNA replication. As expected from prior genomic experiments labeling early replication intermediates with BrdU (Lengronne et al. 2001), we detected strong peaks of EdU incorporation centered on early activating origins of DNA replication along chromosome IV (**Fig 1A**; **middle panel**). The sequences surrounding early origins of replication were enriched ~20-fold relative to late origins (**Supplemental Fig S1A**).

**Figure 1.**
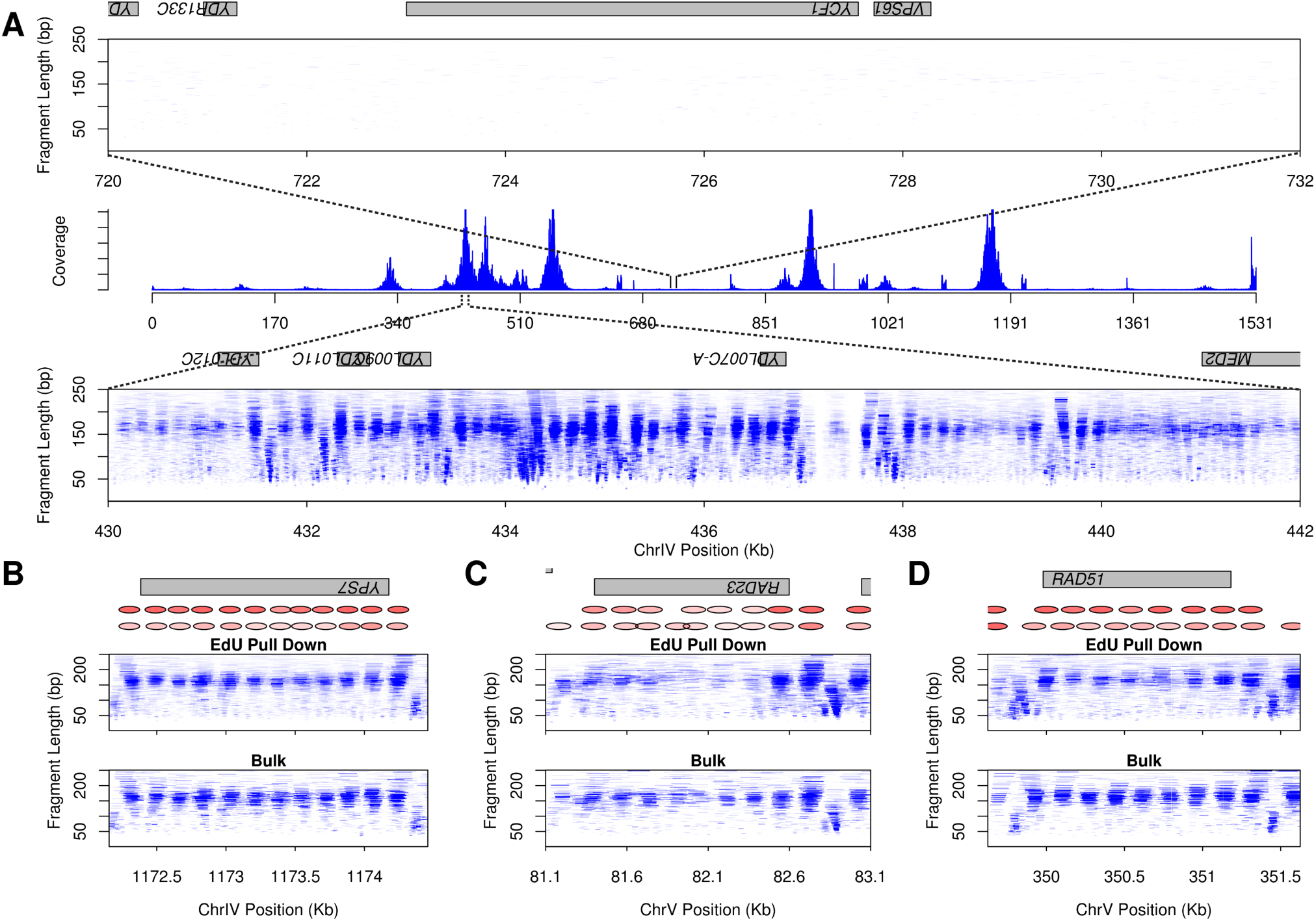
Enrichment of EdU labeled chromatin at sequences proximal to early origins. **A**. Sequencing coverage across chromosome IV. NCOP at an early replicating origin shows a defined chromatin architecture (bottom panel) compared to a non-replicated region (top panel). **B, C** and **D**. Chromatin profiles at genes proximal to an early origin that showed unchanged (**B**) and changed (**C** and **D**) chromatin structure. Nucleosome positions are depicted in red. **D**. RAD51 shows recruitment of a transcription factor and downstream nucleosome shift in the EdU pull down experiment compared to total chromatin.

The innovative aspect of NCOPs is the limited MNase digestion of EdU-enriched chromatin followed by the recovery and sequencing of fragments ~200 bp and smaller. DNA occupancy by a histone octamer will protect a ~150 bp fragment while smaller site-specific DNA binding factors (e.g. transcription factors and replication factors) will typically protect fragments smaller than 80 bp (Henikoff et al. 2011; Belsky et al. 2015). To visualize the NCOPs, we plotted the length of the paired-end reads as a function of their chromosomal position; thus, nucleosomes are evident as well-phased clusters of fragments with lengths of approximately 150 bp and smaller DNA binding factors are evident as discrete clusters of fragments with lengths smaller than 80 bp. We recovered and visualized EdU-labeled chromatin in the vicinity of early activating origins of DNA replication. We found that both nucleosomes and DNA-binding factors were readily distinguishable in the NCOPs from EdU-labeled chromatin (**Fig 1A, bottom**). In contrast, no discernable EdU-enrichment or chromatin organization was detected in the origin distal regions (**Fig 1A, top**), demonstrating the specificity of the NCOP assay for EdU-labeled DNA.

We analyzed the aggregate nucleosome distribution surrounding 539 gene promoters that were within 3500 bp of early origins, and found that nucleosome phasing and occupancy was very similar between the NCOP and ‘bulk’ chromatin (no EdU labeling/enrichment) (**Supplemental Fig S1B**). When examined at the level of individual genes, we found that the recovered EdU-labeled NCOPs largely resemble those prepared from untreated bulk chromatin (**Fig 1B**). However, we did detect a handful of locus-specific alterations in chromatin structure, but these differences were largely attributable to differential transcription of HU-responsive genes. For example, at the *RAD23* locus, we observed displaced nucleosomes from the gene body (**Fig 1C**). At the *RAD51* locus we detected an expansion of the nucleosome free region at the promoter, and the association of an additional DNA-binding factor upstream of the gene in the presence of HU (**Fig 1D**). The recruitment of this factor, which we speculate to be the MBF complex (Leem et al. 1998; Mathiasen and Lisby 2014), was accompanied by a downstream shift in the phasing of the genic nucleosomes. To confirm that these alterations in chromatin structure were due to the HU arrest and not a consequence of the EdU-labeling and enrichment, we also performed an experiment where cells were HU-arrested in the presence of EdU and then released from the arrest for two hours. We found that HU-dependent chromatin changes detected by the NCOPs were restored following release from HU and re-entry into the cell cycle (**Supplemental Fig S1B, right panel, and S1C-S1E)**. Together, these results demonstrate the specificity of NCOPs in detecting EdU labeled chromatin and their sensitivity for detecting local changes in chromatin structure.

### Locus specific differences in the re-establishment of chromatin architecture

Chromatin architecture is disrupted and re-established every cell cycle during the course of S-phase. We sought to survey the dynamics of chromatin maturation throughout the genome by pulse-chase labeling replicating cells with EdU. In order to enrich for cells in S-phase, we first synchronized cells in G1 by the addition of *α*-factor for 2 hours at 24C. The cells were then released into S-phase for 45 minutes and pulsed with EdU for 10 minutes to label nascent chromatin. This was then followed by a 30 minute chase with thymidine to allow the EdU-labeled chromatin to mature (**Fig 2A**). By 45 minutes following α-factor release, the majority of cells were in S-phase with mid-S DNA content (**Supplemental Fig S2A**) allowing us to EdU label replication forks from both early and late firing origins. Consistent with this, we observed a relatively uniform distribution of EdU incorporation across the genome and there were only modest differences in EdU incorporation between early and late replicating sequences (**Supplemental Fig S2B**).

**Figure 2.**
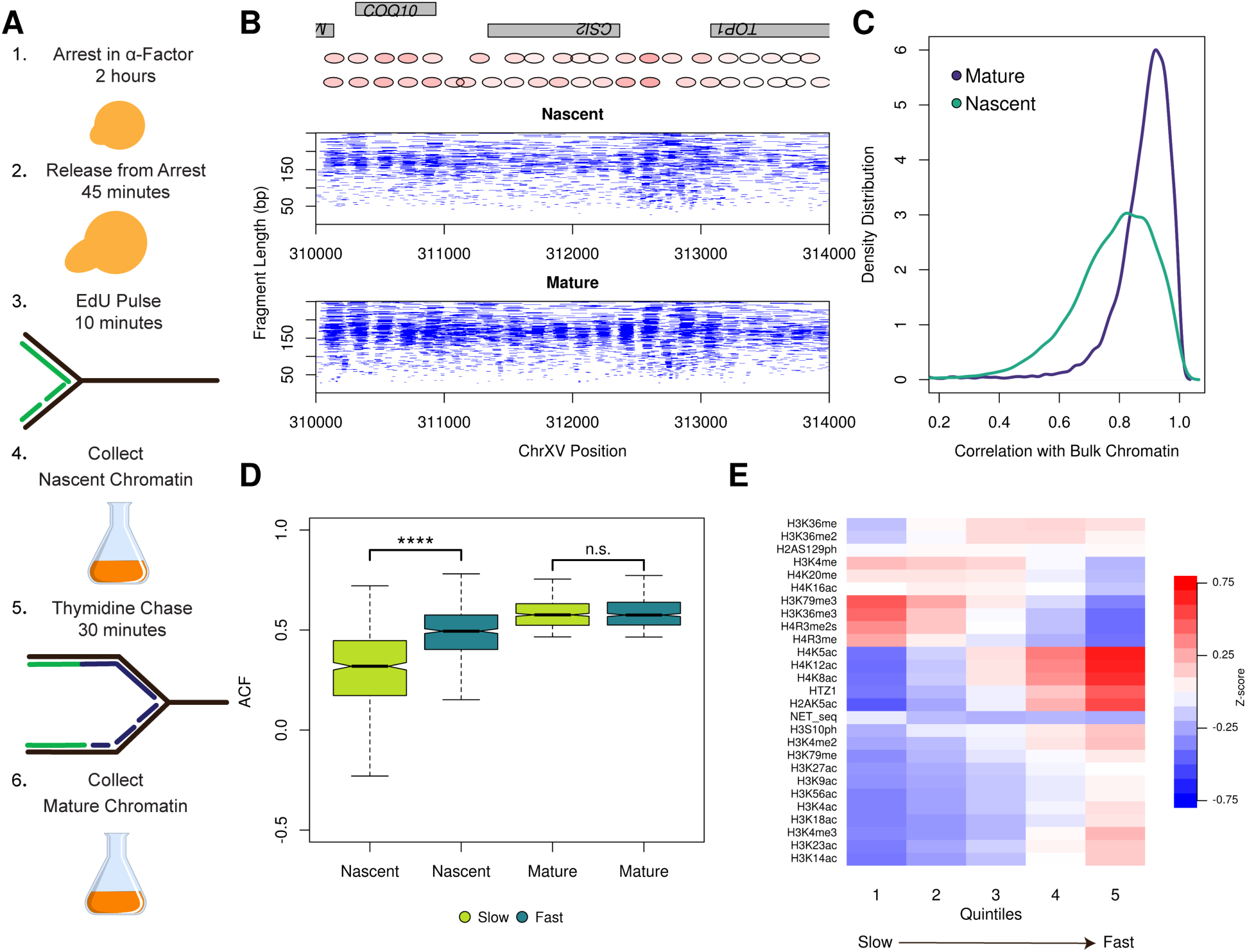
Nascent chromatin occupancy profiles. **A**. Schematic of experimental design for capturing EdU-labeled nascent and mature chromatin. **B**. NCOPs reveal the maturation of chromatin behind the DNA replication fork at a representative locus, *CSI2*. **C**. Nascent chromatin organization in gene bodies is less organized than mature chromatin. Distribution of correlation scores between nascent and bulk (teal) or mature and bulk chromatin (purple). **D**. Differential chromatin maturation kinetics for genes with well-organized mature chromatin. The autocorrelation function (ACF) was used to identify the top 50% of genes with regularly phased arrays of nucleosomes from mature chromatin. These 2700 genes were then binned into quintiles based on the correlation between nascent and mature chromatin. The first and fifth quintiles represent genes with slow and fast chromatin maturation kinetics. The distribution of ACF values (as a proxy for gene organization) in nascent and mature chromatin are depicted for the genes with slow (light green) and fast (dark green) chromatin maturation kinetics. The difference in ACF values among fast and slow maturing chromatin was very significant (p<-22). **E.** Heatmap representing average Z-score values for 25 histone post-translational modifications, the histone variant H2A.Z, and NET-seq scores for the individual quintiles of the correlation between nascent and mature chromatin for the 2700 genes described above.

NCOPs at individual loci revealed that nascent chromatin was inherently more disorganized than mature chromatin. For example, at the *CSI2* locus we observed a marked difference in chromatin organization both upstream and in the gene body between the nascent and mature chromatin samples (**Fig 2B**). To more systematically characterize the differences in nucleosome organization between nascent and mature chromatin, we first used nucleosomal sized reads (140-180 bp) to calculate a nucleosome occupancy profile for each gene in the nascent and mature chromatin state. The nascent and mature nucleosome occupancy profiles were then compared to profiles calculated for bulk chromatin isolated from an asynchronous sample that was not EdU enriched. We found that the structure of mature chromatin at genic regions was highly correlated with bulk chromatin (median R value 0.901); in contrast, nascent chromatin was significantly less correlated (median R value 0.799) and significantly different from the mature chromatin population (p-value 0.73) (**Fig 2C**). We also observed similar patterns from varying the EdU pulse length (5 or 10 min) (**Supplemental Fig S3**). Interestingly, the differences in chromatin organization were not uniform across the genome, but instead seemed to be specified on a gene by gene basis. For example, in contrast to *CSI2*, the chromatin at the neighboring gene *COQ10* was well organized in both the nascent and mature chromatin samples.

To explore the features associated with the maturation dynamics of individual genes, we focused on those genes that exhibited an organized chromatin structure in the mature chromatin state. We used the autocorrelation function (ACF) to determine the regularity of nucleosome phasing and organization within gene bodies, and identified 2700 genes above the median (0.460) of the ACF values. By only focusing on those genes that ultimately have an organized structure in mature chromatin, we were able to explore the range of maturation dynamics from the nascent state. The genes with well organized mature chromatin were stratified into quintiles based on the correlation between nascent and mature chromatin. Because we only focused on those genes with well-organized mature chromatin, their organization in nascent chromatin likely reflects their maturation dynamics with the low (first quintile) and high (fifth quintile) extremes of nascent organization representing slow and fast chromatin maturation dynamics, respectively.

The positioning of the nucleosomes, as revealed by the nucleosome occupancy profiles, was identical between nascent and mature chromatin for those genes in the fifth or fast chromatin assembly quintile. In contrast, the nascent chromatin structure for genes in the first or slow quintile was less organized and readily distinguishable from mature chromatin (**Supplemental Fig S2C**). We then analyzed the distribution of ACF values to examine the regularity of nucleosome phasing and organization within the genes that exhibited fast or slow chromatin maturation dynamics. As expected, the mature state from both classes had a high ACF, indicating well-phased and organized nucleosomes. In contrast, the class of genes with slow chromatin maturation kinetics exhibited significantly poorer nucleosome organization and structure (low ACF) in the nascent as compared to mature state (P-value 1.82e-36) (**Fig 2D**). Together, these data reveal locus-specific differences in chromatin maturation for a large subset of gene bodies.

Our observation that individual genes have distinct maturation dynamics led us to hypothesize that gene-specific chromatin features such as histone post-translational modifications or the occupancy of histone variants are predictive of the slow or rapid chromatin maturation observed at individual genes. To test this hypothesis, we obtained data describing the genome-wide distribution and enrichment of post-translational histone tail modifications and the histone variant H2A.Z (Weiner et al. 2015). We calculated Z-scores of enrichment for each of the chromatin modifications and generated a heatmap of Z-score values for each modification across the quintiles of nascent chromatin organization as described above (**Fig 2E**). We found that genes with rapid chromatin maturation kinetics (fifth quintile) were enriched for H2A.Z and acetylation marks, including those in the lysine residues 5, 8, and 12 of histone H4. Interestingly, H3K4me3 was also enriched in this group of fast maturing genes, and this mark along with H4K12Ac has been shown to colocalize with H2A.Z (Chen et al. 2012). There was also a significant decrease in the chromatin marks associated with actively transcribed genes, including H3K36me3 and H3K79me3. Consistent with the relative depletion of marks associated with active transcription, we found that these genes with rapid chromatin maturation exhibited less active transcription as determined by native elongating transcript sequencing (NET-seq) data (Churchman and Weissman 2011). In contrast, we observed a depletion of the H2A.Z variant and enrichment of transcription marks in the genes with slow maturation kinetics (first quintile).

### Nucleosomes exhibit distinct patterns of positioning and occupancy genome wide

The yeast genome is relatively gene dense compared to metazoan genomes; however, 40% of the genome remains intergenic. Thus, while our prior analysis focused on chromatin maturation in gene bodies, we also sought to explore chromatin maturation at the level of individual nucleosomes. We first identified the position of the nucleosome dyad for ~70,000 high confidence nucleosomes from bulk chromatin. For every nucleosome sized fragment recovered from either the nascent or mature chromatin, we calculated the distance of the fragment midpoint to the nearest nucleosome dyad in bulk chromatin. The average of these distances for each nucleosome represented a positioning score, with well-positioned nucleosomes having a low score, and poorly positioned nucleosomes having a high score. Genome-wide, we found that individual nucleosomes in nascent chromatin are more poorly positioned as compared to mature or bulk chromatin (**Fig 3A**).

**Figure 3.**
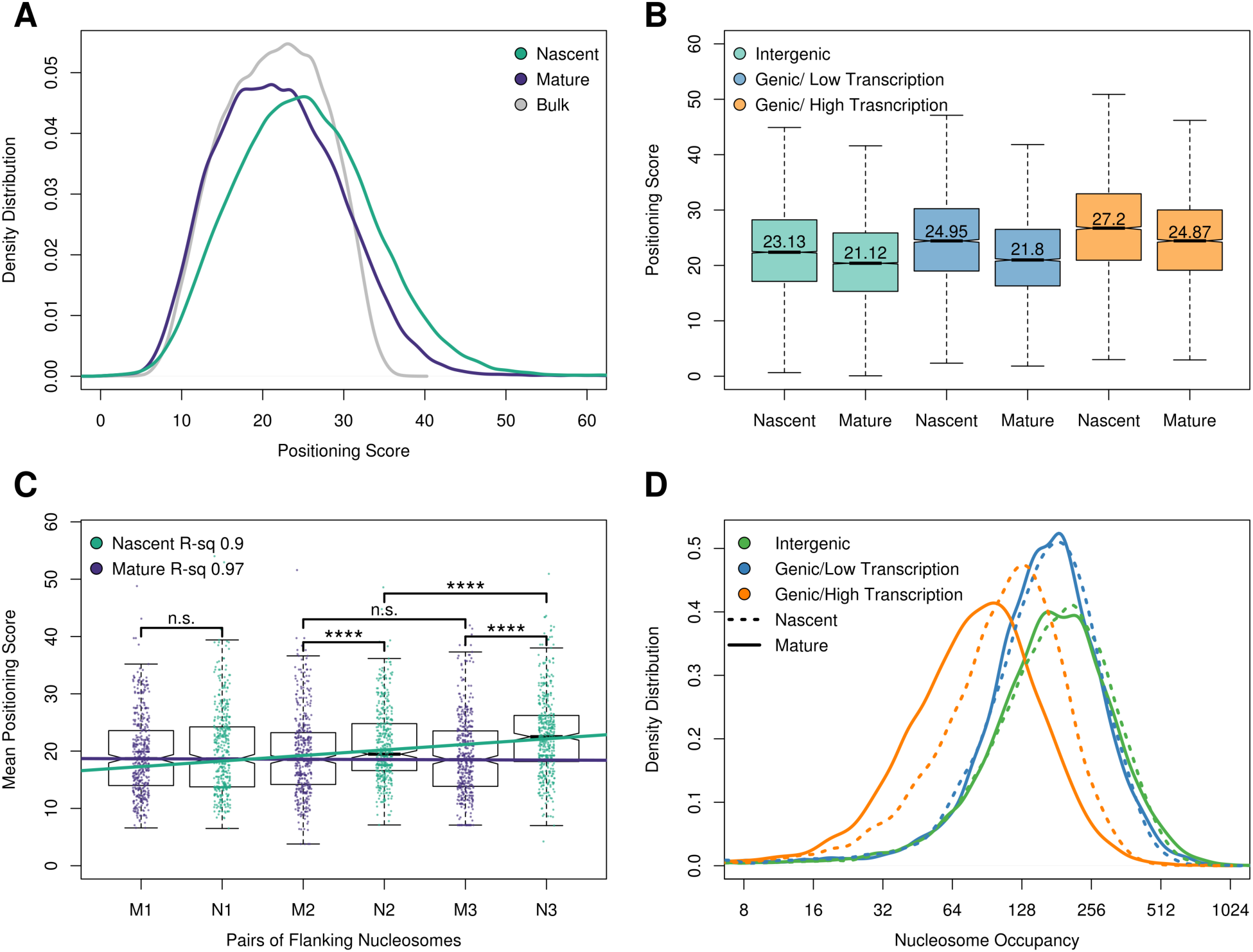
Genome-wide nucleosome positioning and occupancy in nascent and mature chromatin. **A**. Genome-wide distribution of nascent, mature and bulk nucleosome positioning. ~70,000 high-confidence nucleosome dyads were identified in bulk chromatin. For each chromatin fraction (nascent, mature and bulk) the distance from the midpoint of each sequencing read to the nearest nucleosome dyad was calculated as a nucleosome positioning score. **B**. Nascent and mature chromatin organization (nucleosome positioning scores) at intergenic and intragenic nucleosomes. Intragenic nucleosomes were subdivided into high (top 10%) and low (bottom 10%) transcriptional activity. **C**. Transcription factors influence chromatin maturation kinetics. Positioning scores of the first, second, and third pairs of nucleosomes flanking 436 transcription factors with strong occupancy in mature chromatin. *T*-test **** p ≤ 0.0001; n.s. p > 0.05. **D**. Active transcription displaces nucleosomes from mature chromatin. Distribution of nucleosome occupancy scores (sequencing reads assigned to individual nucleosomes) for nascent and mature chromatin at intragenic and intergenic sequences. Actively transcribed genes have more nucleosome occupancy in nascent chromatin than in mature chromatin (P-value 4.80 x 10^-67^).

Individual nucleosomes were broadly classified as either intergenic or genic. We found that nucleosomes within intergenic regions have better positioning scores than nucleosomes within gene bodies. To begin to understand the differences in chromatin maturation for intergenic and genic nucleosomes and their relationship to transcription, we identified the nucleosomes associated with the most (top 10%) and least (bottom 10%) expressed genes as determined by prior NET-Seq experiments (Churchman and Weissman 2011) (**Fig 3B**). A similar number of nucleosomes were sampled from intergenic regions. In each category, we found that the positioning of the nucleosomes was decreased in nascent relative to mature chromatin. Nucleosomes in intergenic and poorly transcribed genes were better positioned than nucleosomes from active genes, consistent with transcription dependent nucleosome eviction and remodeling (Lee et al. 2004; Boeger et al. 2004; Bernstein et al. 2004).

Transcription factors such as Abf1p and Rap1p are thought to play a critical role in establishing nucleosome organization and chromatin structure (Yarragudi et al. 2004; Ganapathi et al. 2011). We reasoned that the greater nucleosome organization observed in intergenic regions was due, in part, to the presence of transcription factors. To assess the chromatin maturation dynamics at intergenic regions, we examined nucleosome positioning for the first, second and third pairs of nucleosomes surrounding 436 predicted transcription factor binding sites (MacIsaac et al. 2006) with defined occupancy footprints in mature chromatin (**Fig 3C**). We found no significant difference in nucleosome positioning for the mature nucleosomes. In contrast, we identified a significant distance-dependent increase in nucleosome positioning scores for each successive nucleosome pair in the nascent chromatin. Together, these results underscore the role of transcription factors functioning as barrier elements in establishing nucleosome organization (Zhang et al. 2009) following DNA replication.

To further explore the dynamics of chromatin maturation, we also examined nucleosome occupancy for intergenic and genic regions of the genome. An occupancy score was calculated for individual nascent and mature nucleosomes as the number of fragment midpoints mapping within 70 bp of the high-confidence nucleosome dyads identified from bulk chromatin (see above). We found that nucleosomes in intergenic regions and genes that were not being actively transcribed exhibited similar nucleosome occupancies in both nascent and mature chromatin states (**Fig 3D**). Thus, while nucleosomes are rapidly deposited behind the replication fork, they do not converge on their preferred position until maturation. In contrast to intergenic and non-transcribed regions, we observed significantly more nucleosome occupancy in nascent relative to mature chromatin in actively transcribed genes, suggesting that the newly deposited nucleosomes behind the DNA replication fork are evicted by active transcription.

### Actively and passively replicated origins have distinct maturation dynamics

Replication forks must traverse the entire genome during S-phase but emanate from a select few DNA replication origins. Each licensed origin has its own inherent efficiency of activation during S-phase (Aparicio 2013; Hawkins et al. 2013). A consequence of this is that highly efficient origins will initiate bidirectional DNA replication every cell cycle whereas the least efficient origins will be passively replicated by forks from neighboring efficient origins. We reasoned that there might be distinct differences in chromatin maturation at active origins that initiate DNA replication versus those origins that are passively replicated. The distribution and strandedness of Okazaki fragments around origins were used as a proxy for origin efficiency (McGuffee, Smith, and Whitehouse 2013). We first examined locus specific NCOPs for *ARS805* and *ARS602*, an efficient early origin and a passively replicating origin, respectively (**Fig 4A-B**). As previously reported, we observed well-ordered and phased nucleosomes flanking either origin in mature chromatin (Belsky et al. 2015). We also detected smaller fragments (< 80 bp) in the mature chromatin that are indicative of ORC binding in the NFR at the ARS consensus sequence of both origins. In contrast, we observed a difference in nascent chromatin organization between *ARS805* and *ARS602*. Specifically, we found that there was significantly more nascent chromatin organization at the active replication origin as compared to the passively replicated origin.

**Figure 4.**
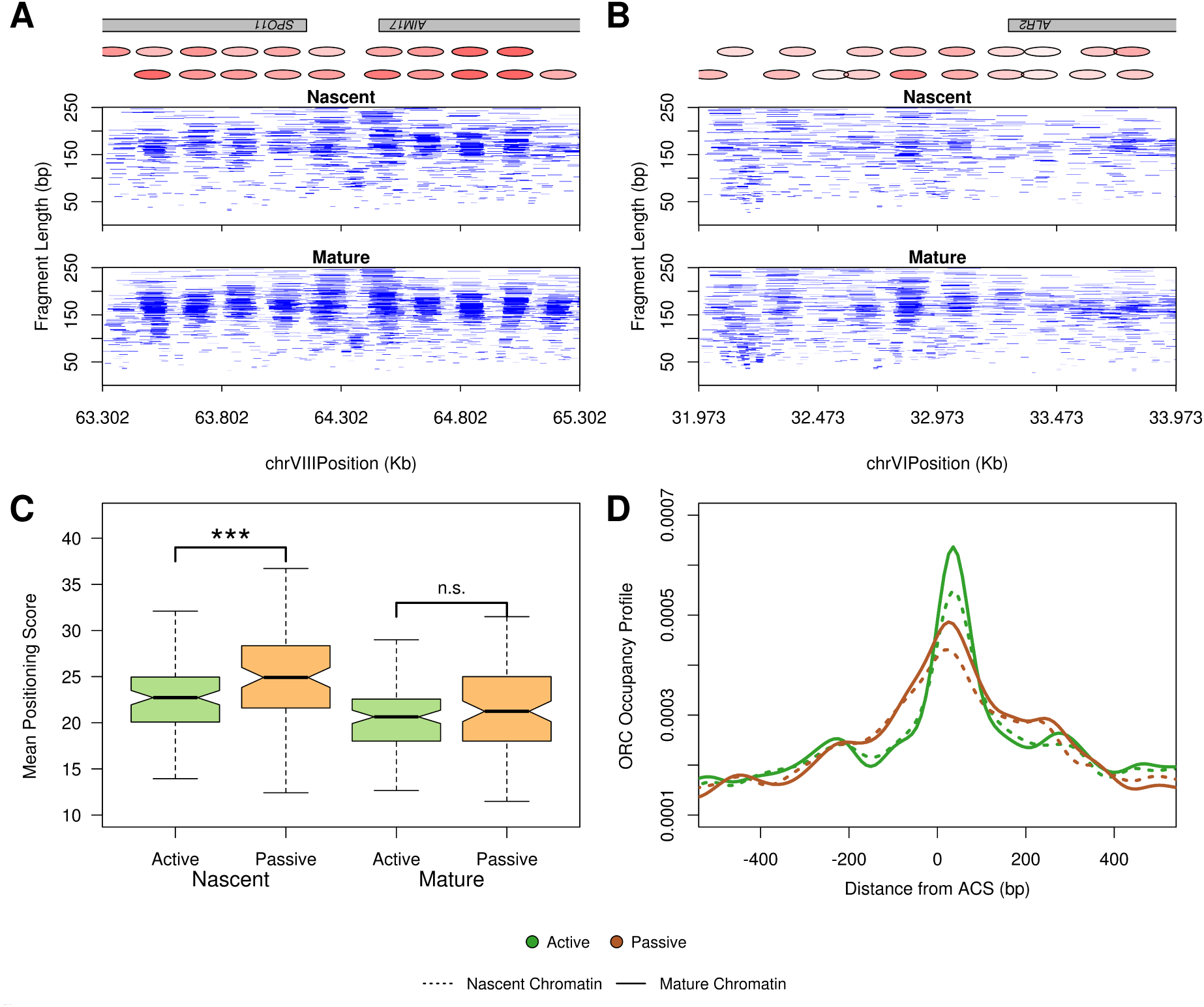
Chromatin maturation at actively and passively replicated origins. **A.** Chromatin occupancy profile for nascent and mature chromatin at the active origin *ARS805*. **B**. Chromatin occupancy profile for nascent and mature chromatin at the inactive origin *ARS602*. **C**. Passively replicated origins have slower chromatin maturation kinetics. Boxplots depicting the distribution of nucleosome positioning scores for nascent and mature chromatin at 100 active and 100 inactive origins. Nascent chromatin surrounding passively replicated origins is more disorganized than at active origins. *T*-test *** p ≤ 0.001; n.s. p > 0.05. **D**. ORC occupancy is decreased at passively replicated origins. Density distribution of ORC occupancy footprints as determined from the NCOPs.

To more comprehensively examine chromatin maturation dynamics at active and passive origins, we identified 261 origins with an ORC-dependent footprint (Siow et al. 2012; Belsky et al. 2015) and then stratified the origins by their efficiency score to identify the top 100 active and passive replication origins. The mean efficiency for each class of origins was 0.65 (active) versus 0.017 (passive). For each actively or passively replicated origin, we examined the nucleosome positioning scores for the first three nucleosomes flanking each origin up and downstream (**Fig 4C**), where increasing chromatin organization is represented by lower nucleosome positioning scores. We found that the nascent chromatin surrounding passive origins was significantly more disorganized than at active origins. In contrast, there were no detectable differences in chromatin organization between active and passive origins in the chromatin that had matured behind the replication fork. Our NCOPs also revealed a decrease in small fragments (<80 bp) at the passively replicated origins, consistent with decreased ORC occupancy (**Fig4D**). Together, these results suggest a feedback mechanism in place at active origins to promote ORC recruitment and immediately re-establish chromatin architecture for the next cell cycle.

### Transient association of DNA binding factors with nascent chromatin

DNA binding proteins, such as transcription factors and ORC, associate with specific primary sequences; however, for any given factor there are many more potential sequence motifs than occupied sites in the genome (Breier, Chatterji, and Cozzarelli 2004; Eaton et al. 2010; Wang et al. 2012; Slattery et al. 2014). Nucleosome occupancy is thought to limit access to many of these potential motif matches, thus defining the regulatory landscape. NCOPs provided an opportunity to comprehensively survey DNA occupancy in a factor agnostic manner throughout the genome in both nascent and mature chromatin. We hypothesized that as nucleosomes are deposited and become organized in nascent chromatin, there may be transient or promiscuous interactions between transcription factors and DNA immediately behind the replication fork.

To identify all potential sites of DNA occupancy that were not protected by a nucleosome, we focused on the paired-end sequencing fragments that were less than 80 bp and identified 6272 loci that were significantly enriched (*p*<0.05) for these size fragments in both nascent and mature chromatin. To evaluate changes in DNA-binding factor occupancy, we first determined the log2 ratio of normalized occupancy scores for nascent and mature chromatin at each of the 6272 sites and plotted these values as an ordered heatmap (**Fig 5A**). In general, the heatmap revealed three classes of DNA binding profiles which were indicative of their chromatin maturation dynamics: i) slow maturation -- sites with greater occupancy in mature chromatin; ii) fast maturation -- sites that were equally occupied in both mature and nascent chromatin; and iii) transient occupancy -- sites that were enriched in nascent and not mature chromatin.

**Figure 5.**
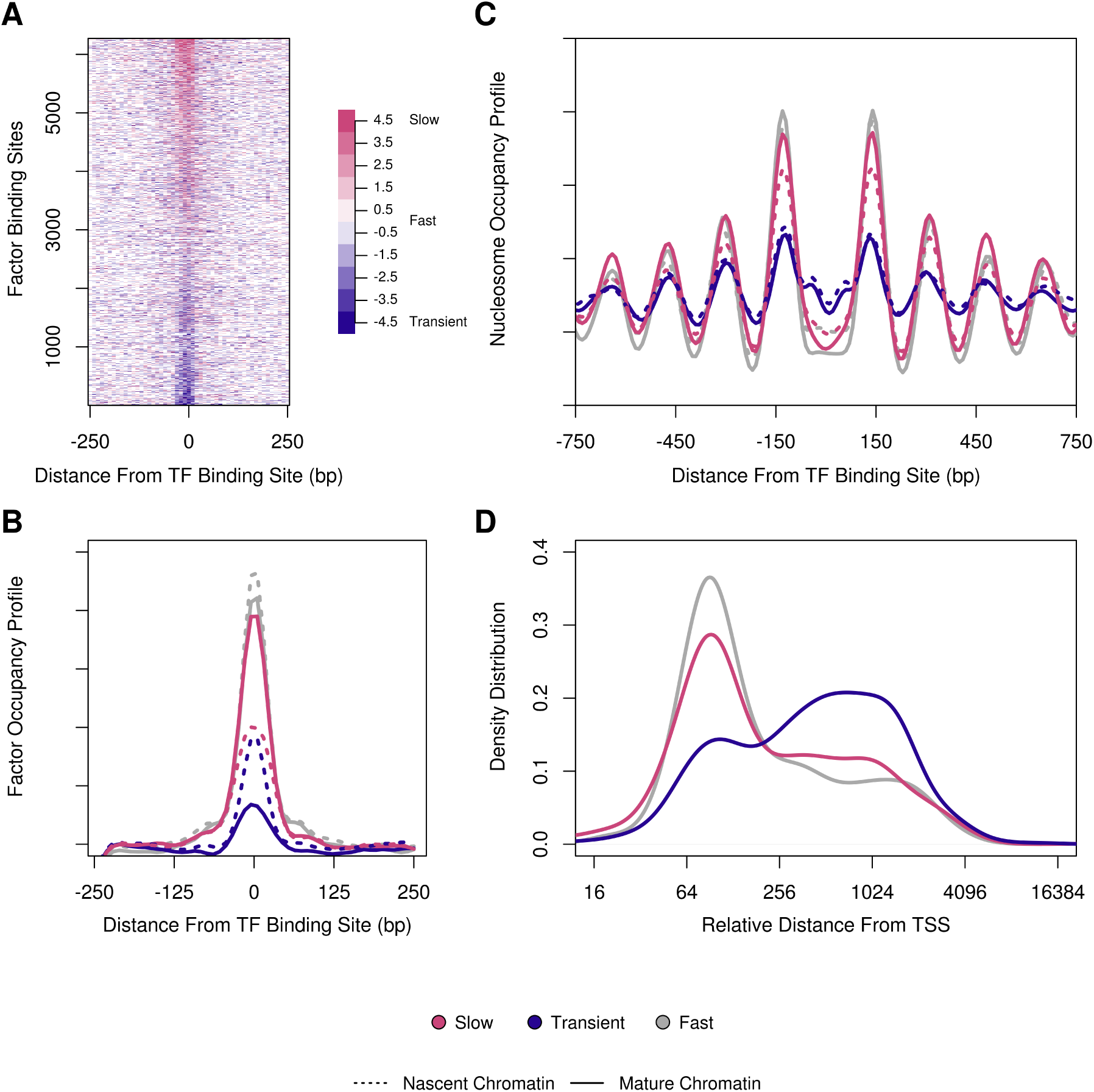
Transcription factor association with nascent and mature chromatin. **A**. Ratio heatmap log2(mature/nascent) of 6272 high-confidence sites of small DNA binding factor occupancy obtained from DNA fragment lengths between 20 bp and 80 bp in length. **B.** Identification of three classes of chromatin maturation for DNA binding factor occupancy: fast -- equal occupancy in nascent and mature chromatin; slow -- greater occupancy in mature than nascent chromatin; and transient -- greater in nascent than mature. **C**. Average plot of nucleosome organization at sites occupied by the factors described in (B). **D**. Sites of transient occupancy are promiscuously enriched at sites distal from the transcription start site (P-value 2.01 x 10^-19^ between fast and transiently associating factors).

To better characterize the chromatin maturation of regulatory sites bound by transcription factors and other DNA-binding factors, we identified the extreme deciles of sites (~625 sites for each decile) representing locations with slow or transient occupancy and an equal number of loci with fast maturing kinetics (**Fig 5B**). As expected, the occupancy of small DNA-binding factors in the sites with slow maturation kinetics was less in nascent than mature chromatin (compare dashed and solid red lines). Despite the different kinetics of maturation, similar occupancy levels were ultimately established in the mature chromatin (compare solid red and grey lines).

We analyzed the nucleosome organization surrounding slow, fast, and transiently maturing sites of DNA-binding factor occupancy (**Fig 5C**). We found that both the slow and fast maturing sites of DNA occupancy were surrounded by well positioned nucleosomes in both nascent and mature chromatin. This suggests that for at least some sites, defined nucleosome positioning may be a requisite for factor occupancy. In contrast, we found that specific pioneer factors (e.g. Abf1p and Reb1p) exhibited occupancy footprints and chromatin organization profiles that were indistinguishable between nascent and mature chromatin, consistent with their immediate deposition behind the DNA replication fork (**Supplemental Fig S4**).

The sites with transient occupancy only observed in the nascent chromatin suggest that they may represent promiscuous binding sites to accessible motifs in regions of poor chromatin organization. The compact nature of the yeast genome dictates that most regulatory DNA binding sites are proximal to gene promoters. Thus, we examined the location of each binding site for all three occupancy classes -- fast, slow and transient -- relative to their gene start site. The DNA binding sites with fast and slow maturation were found proximal to promoters (**Fig 5D**). In contrast, the distribution of transient sites was not limited to promoters, but rather occurred distally to promoters and likely in gene bodies. Together, these results suggest that potential binding motifs are exposed in poorly organized nascent chromatin and that they are ultimately removed by chromatin maturation or transcription.

## Discussion

In proliferating cells, it is critical that both genetic and epigenetic information be faithfully transmitted between generations. One of the consequences of semi-conservative DNA replication is that the chromatin landscape needs to be disrupted ahead of the replication fork and then re-established in its wake. While genetic and biochemical experiments have elucidated many of the factors and mechanisms regulating the inheritance and assembly of chromatin behind the fork (Smith and Stillman 1989; Shibahara and Stillman 1999; Tyler et al. 1999; Q. Li et al. 2008; Sirbu et al. 2013; Alabert et al. 2014), the spatio-temporal dynamics of chromatin assembly across the genome are just starting to be revealed (Fennessy and Owen-Hughes 2016; Ramachandran and Henikoff 2016; Vasseur et al. 2016). We have utilized nascent chromatin occupancy profiles (NCOPs), which combine MNase epigenome mapping with EdU-labeling of nascent DNA to provide a factor agnostic view of chromatin maturation at nucleotide resolution across the *S. cerevisiae* genome.

The assembly of chromatin during DNA replication is a rapid process facilitated by replication-dependent histone chaperones and assembly factors (Alabert and Groth 2012; Serra-Cardona and Zhang 2018). Despite the rapid deposition of histone octamers behind the replication fork, nascent chromatin exhibits increased nuclease sensitivity (DePamphilis and Wassarman 1980; Klempnauer et al. 1980; Annunziato and Seale 1982; Stillman 1986) suggesting poorly organized nucleosomes relative to mature chromatin. The establishment of chromatin organization is thought to be dependent on primary sequence (Segal et al. 2006; N. Kaplan et al. 2009), transcription (Weiner et al. 2010), and the positioning of nucleosomes relative to fixed barrier elements (Bai et al. 2010; M. Li et al. 2015; Yan, Chen, and Bai 2018). In addition, our NCOPs revealed locus-specific differences in chromatin maturation kinetics at genes and origins of replication suggesting that the maturation process is also influenced by the local chromatin state and origin activity.

We first focused on the chromatin maturation dynamics of genes with well-organized nucleosome architecture in mature chromatin. Among the genes with well-positioned nucleosomes in mature chromatin, we were able to rank them by their nascent chromatin organization and identify those genes that had either poorly organized (slow maturing) or well-organized (fast maturing) chromatin organization. In contrast to prior reports describing transcription-dependent chromatin maturation dynamics in yeast (Vasseur et al. 2016), we found that maturation of nascent chromatin was fastest for genes with low transcriptional activity (**Fig 2E**). The Vasseur study focused on the temporal maturation of nascent chromatin using the correlation with mature chromatin as a proxy for maturation within the gene body. As active transcription frequently results in nucleosome eviction and disorganized phasing in bulk chromatin (Lee et al. 2004; Boeger et al. 2004; Bernstein et al. 2004), they were looking at a de facto transcription-dependent effect. We also note in our studies that we detected the transcription-dependent eviction of nascent histone octamers from actively transcribed gene bodies (**Fig 3D**) consistent with the role of transcription in shaping the mature chromatin landscape.

A long standing model for how nucleosome positioning is achieved proposes that barrier elements such as transcription factors aid in establishing the fixed positions of their flanking nucleosomes (Fedor, Lue, and Kornberg 1988; Pazin et al. 1997; Mavrich et al. 2008; M. Li et al. 2015). As a result, this poses further positioning constraints on the subsequent +2 and +3 nucleosomes which thereby stabilizes their positioning. This is because the nucleosomes are packaged within short genomic regions, and the fixed position of one nucleosome influences the organization of adjacent ones in the array. General regulatory factors (GRFs) like Abf1p and Reb1p are capable of positioning flanking nucleosomes without the need for chromatin remodeling (Yarragudi et al. 2004; Hartley and Madhani 2009; Ganapathi et al. 2011), and can position nascent nucleosomes following passage of the replication fork (Yadav and Whitehouse 2016). In agreement with this, we find that the first pair of nucleosomes immediately flanking factors bound to mature chromatin are tightly positioned in both nascent and mature chromatin (**Fig 3C**), indicating that the association of regulatory factors with DNA is important to position nucleosomes behind the fork.

Histone post-translational modifications and histone variants have long been postulated to form the basis of epigenetic memory (Probst, Dunleavy, and Almouzni 2009; MacAlpine and Almouzni 2013). The local inheritance of parental H3/H4 histones at the replication fork, in part, contributes to the re-establishment of the parental chromatin state on the newly replicated DNA. As the chromatin maturation dynamics we observed appeared to be specified on a gene by gene basis, we hypothesized that the local parental chromatin state may influence chromatin maturation dynamics on the nascent DNA. To explore this idea, we systematically examined a comprehensive panel of histone post translational modifications and histone variants derived from an asynchronous yeast population (**Fig 2E**) (Weiner et al. 2015). We found that genes with rapid maturation kinetics were poorly expressed and depleted of modifications frequently associated with gene expression (e.g. H3K36me3 and H3K79me3) and instead were enriched for specific histone modifications including H4K12ac, H3K4me3 and the histone variant H2A.Z. In *S. cerevisiae,* H2A.Z is located at the promoters of inactive or poorly transcribed genes, and helps to stabilize their promoter architecture for recruitment of regulatory factors (Guillemette et al. 2005; B. Li et al. 2005). Deposition of parental H3/H4 tetramers containing H4K12ac and or H3K4me3 behind the fork may help facilitate the recruitment of H2A.Z (htz1p) to inactive promoters, as single molecule studies revealed that Htz1 variant containing nucleosomes are enriched for H3K4me3 and H4K12Ac (Chen et al. 2012). The recruitment of H2A.Z may rapidly stabilize nascent nucleosome organization by inhibiting ATP-dependent chromatin remodeling activity (B. Li et al. 2005).

We also investigated the chromatin maturation dynamics surrounding other chromosomal features including origins of DNA replication. The timing and efficiency by which individual origins of DNA replication are activated during S-phase is determined, in part, by primary sequence, rate limiting initiation factors (Mantiero et al. 2011; Tanaka et al. 2011), and the local chromatin structure (Berbenetz, Nislow, and Brown 2010; Eaton et al. 2010; Kurat et al. 2017). We found that the maturation dynamics of the chromatin surrounding origins of DNA replication was dependent on whether the origin actively initiated DNA replication or was passively replicated by a replication fork from elsewhere in the genome. Efficient origins that initiate replication in the majority of cell cycles (McGuffee, Smith, and Whitehouse 2013) rapidly reset their chromatin structure and exhibit well positioned nucleosomes flanking the origin immediately after initiation in nascent chromatin. The differential maturation of chromatin between efficient and passive origins likely reflects distinct molecular mechanisms in the re-assembly of chromatin following either an initiation event or the disassembly of the pre-replication complex at inefficient origins. It is tempting to speculate that immediately following unwinding of the origin DNA and initiation of DNA replication, an active mechanism promotes the rapid re-association of ORC and precise phasing of nucleosomes which have become a hallmark for eukaryotic origins (Berbenetz, Nislow, and Brown 2010; Eaton et al. 2010; Lubelsky et al. 2011; Cayrou et al. 2015; Miotto, Ji, and Struhl 2016).

Chromatin maturation is governed by both replication-dependent and independent processes. While the deposition of nascent histone octamers behind the fork is replication-dependent, the re-establishment of the regulatory landscape and binding of transcription factors is dependent on chromatin remodeling, local epigenetic signatures and recruitment of other trans-acting factors (Allis and Jenuwein 2016). For most transcription factors, there are many more potential motif matches in the genome than there are sites occupied by the factor. Competition between nucleosome occupancy and transcription factor binding has long been thought to be a determinant of which regulatory motifs are occupied (Wasson and Hartemink 2009; M. Li et al. 2015; Ramachandran and Henikoff 2016). The factor agnostic NCOPs provided insight into how trans-acting factors and the regulatory landscape was established during chromatin maturation. Not surprisingly, we found a range of maturation kinetics for loci occupied by non-nucleosomal DNA binding factors, including those that rapidly associate with nascent chromatin (fast) and those that associate later in mature chromatin (slow). However, we also identified a significant number of sites that were only transiently occupied in nascent chromatin. Unlike the binding sites with slow or fast kinetics which were promoter proximal, the transient sites were frequently located distal to the promoter and in gene bodies. This is consistent with a promiscuous or opportunistic mode of binding in the absence of organized chromatin immediately following DNA replication (Yan, Chen, and Bai 2018). During development in higher eukaryotes, the DNA replication program is characterized by changes in the number of firing origins, the length of S-phase, and the timing of replication (Duronio 2012; Rhind and Gilbert 2013). The developmental plasticity in the DNA replication program may lead to promiscuous binding of regulatory factors through the chromatin changes that occur throughout this process, and thus contribute to epigenetic regulation and cell-type specific gene expression programs.

## Acknowledgements

We thank members of the MacAlpine lab for critical comments and suggestions. We also thank Jason Belsky for advice on data analysis. This work was supported by the National Institutes of Health R35 GM127062 DMM and F31-GM115158 MPG.

## Methods

### Yeast strains

The yeast strain DMMy218 is in the W303 background and has the genotype *MATa*, *leu2-3,112*, *BAR1::TRP*, *can1-100*, *URA3::BrdU-Inc*, *ade2-1, his3-11,15*.

### Chromatin occupancy profiling

To label cells in early S-phase, yeast were grown in rich medium at 30C to an O.D of ~0.7 and arrested in G1 phase with α-factor (GenWay) at a final concentration of 50 ng/mL for 2 hours. Cells were then washed twice in sterile water, resuspended in fresh medium containing 0.2 M hydroxyurea (Sigma) and 130 µM EdU (Berry & Associates, Inc), and grown for 2 hours. Cells were washed twice with sterile water and the pellets were quick-frozen and stored at −80C.

In order to profile nascent and mature chromatin, yeast cells were grown at 25C to an O.D of ~0.7 and arrested in G1 phase with α-factor at a final concentration of 50 ng/mL for 2 hours. Cells were then washed twice in sterile water, resuspended in fresh medium and allowed to grow for 45 minutes to enter S phase. EdU was then added to a final concentration of 130 µM and allowed to grow for 10 minutes (pulse), after which a sample was taken for nascent chromatin. Cells were washed, resuspended in fresh medium containing 1.3 mM thymidine (Sigma), and allowed to grow for 30 minutes (chase) when a sample was taken for mature chromatin. Cells were washed twice with sterile water and the pellets were quick-frozen and stored at −80C. All experiments were performed using independent biological replicates.

### Chromatin preparation

MNase digestions were performed as described in (Belsky et al. 2015).

### Click reaction and streptavidin affinity capture

100 µg of MNase-digested DNA were concentrated in a speed vacuum. DNA was incubated in click chemistry reaction buffer as described in (Kliszczak et al. 2011; Sirbu, Couch, and Cortez 2012; Leung, Abou El Hassan, and Bremner 2013). CuSO4 and ascorbic acid were replaced with CuBr and TBTA (Sigma). The click reaction proceeded for 1 hour at room temperature with gentle shaking. DNA was recovered by ethanol precipitation and resuspended in 30 µl sterile water.

Biotin conjugated EdU-labeled DNA was enriched using 5 µl streptavidin magnetic beads (New England Biolabs). Beads were resuspended in blocking solution (2 % I-Block (Thermofisher), 25 mM Tris, 150 mM NaCl, and 0.5 % SDS) for 2 hours with gentle shaking at room temperature. Beads were then washed twice with cold binding buffer (Leung, Abou El Hassan, and Bremner 2013). Recovered DNA was added to blocked beads and incubated in 200µl cold binding buffer for 1 hour at 4°C. Bead-bound DNA was washed once with binding buffer followed by 3 washes with EB buffer (Qiagen).

### Flow cytometry

To analyze yeast cells by flow cytometry, cells were resuspended in 70% ethanol and fixed overnight at 4C. Then, cells were washed, sonicated, and incubated in 50 mM sodium citrate pH7.4 with 0.3 mg/ml RNase A for 2 hours at 50C. 0.6 mg/ml proteinase K was then added and incubated for an additional 2 hours at 50C. Finally, cell pellets were resuspended in 50 mM sodium citrate and 1:5000 sytox green (Invitrogen) and incubated for 1 hour at room temperature. Flow cytometry was performed on a BD FACSCanto analyzer and 30,000 cells were recorded for each sample.

### Sequencing library preparation

Illumina sequencing libraries were prepared as previously described (Henikoff et al. 2011; Belsky et al. 2015), with the following modifications: All library preparations were done on bead-bound DNA. After each step, clean up was accomplished by washing beads twice with binding buffer and three times with EB buffer. NEBNext Multiplex Oligos for Illumina kit (New England Biolabs) was used in adapter ligation and PCR steps. PCR reactions were cleaned using Agencourt AMPure XP beads (Beckman).

### Data Analysis

Data analysis was performed in R version 3.2.0 or using bash scripts. Data processing scripts are available at https://gitlab.oit.duke.edu/dmm29/Gutierrez_2018

### Sequence Alignment

Sequencing reads were aligned in paired-end mode to the sacCer3/R61 version of the *S. cerevisiae* genome using Bowtie 0.12.7 (Langmead et al. 2009).

### Analysis of chromatin organization and structure

Biological replicates were merged for all data analyses. Merged files were then sampled to obtain equal read depth. Each merged experiment represents a minimum of 30 million reads. To characterize chromatin structure in nascent and mature chromatin at gene bodies, nucleosome-size reads between 140 bp and 180 bp in length were used to calculate a density curve of nascent, mature, and bulk chromatin across the length of each gene, with a 30 bp bandwidth Gaussian kernel. The pairwise pearson correlation between the density curves at each gene was calculated for nascent and bulk, or mature and bulk chromatin.

Similarly, we obtained nucleosome-size reads overlapping each gene and determined the midpoint location of each. We then determined the density distribution of all the read midpoints with a 30 bp bandwidth Gaussian kernel. This output signal was smoothed and the refined predicted values generated a curve where each peak represents a nucleosome dyad and each trough represents linker DNA. We used an autocorrelation function (ACF) to determine if the pattern of positioning for the first four nucleosomes within each gene changed or remained organized. This ACF value was used as a proxy for individual gene structure.

### Signal normalization of histone post-translational modifications, H2A.Z variant occupancy, and NET-seq

The accession number GSE61888 was used to locate and download ChIP-seq raw histone post translational modification (PTM) and htz1 histone variant data from (Weiner et al. 2015). Raw NET-seq data was obtained using the accession number GSE25107 (Churchman and Weissman 2011). Reads were aligned in single end mode to the sacCer3/R61 version of the *S. cerevisiae* genome using Bowtie 0.12.7 (Langmead et al. 2009).

To determine PTM and htz1 occupancy, we calculated all the mapped reads overlapping gene bodies and normalized read counts using RPKM (Reads Per Kilobase of transcript per Million mapped reads). NET-seq data was processed similarly; however, the enrichment of reads was determined for sequences that spanned 100 bp upstream of the gene start to 300 bp downstream of the gene start. Gene orientation was considered during this calculation.

### Individual nucleosome positioning and occupancy scores

Nucleosome locations were determined by obtaining nucleosome-size reads between 140bp and 180 bp in length and calculating a density curve across the length of each chromosome with a 30 bp bandwidth Gaussian kernel. Each chromosome-length density calculation was passed through the turnpoints function in the pastecs R package to determine all the potential peaks from the density curves. These peaks were used to calculate nucleosome coordinates 70 bp upstream and downstream from each peak (140 bp total width).

All the nucleosome-size reads from each chromosome were acquired and their midpoint positions determined. Then, the distance of each midpoint to its nucleosome dyad was calculated. This average was used as the nucleosome positioning score. To ascertain nucleosome occupancy, the total number of normalized sequencing reads overlapping a nucleosome was determined.

### Data Availability

All genomic data is publicly available at NCBI SRA accession SRP158706

## Supplemental Figure Legends

**Supplemental Figure S1.**
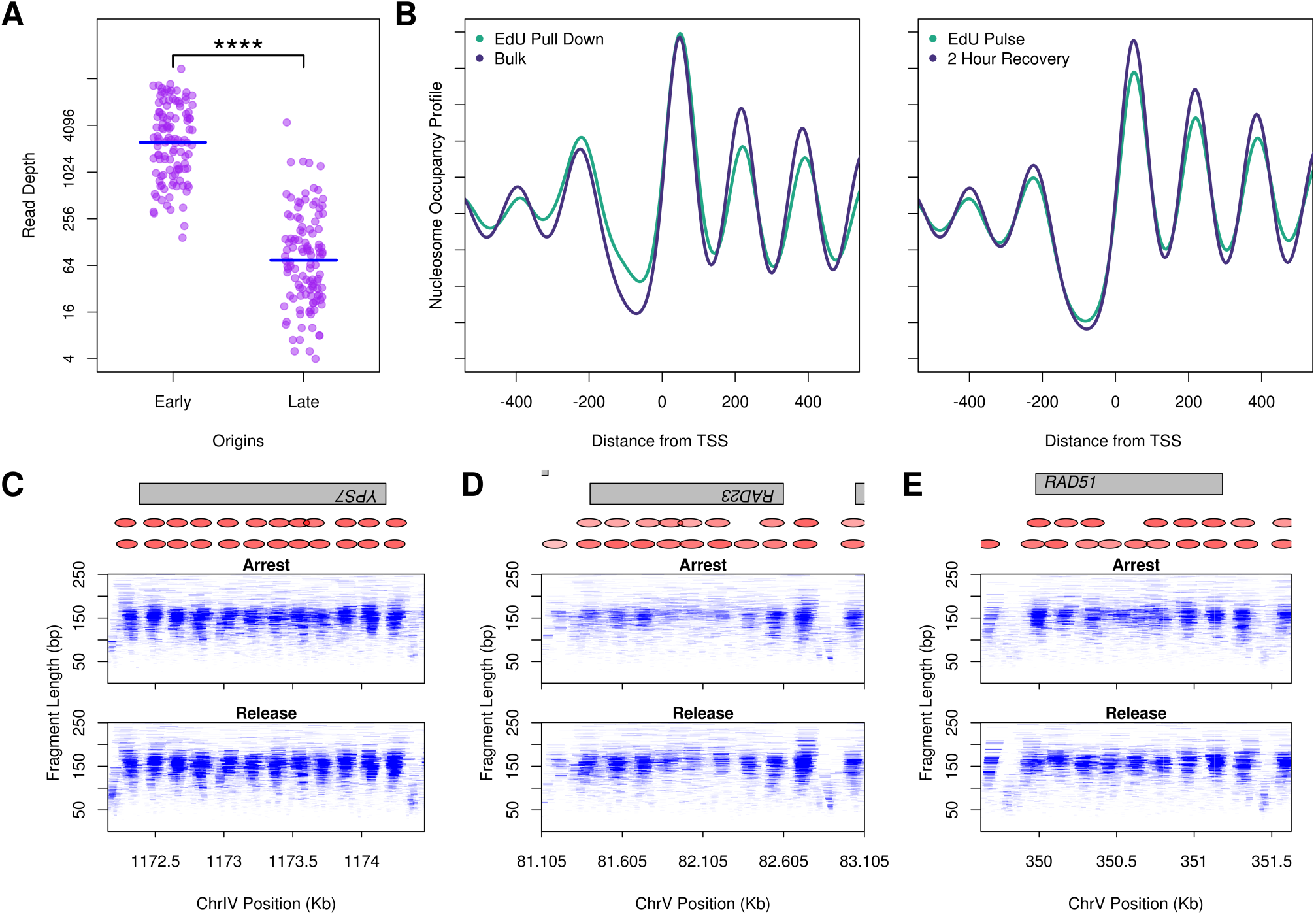
EdU-labeling and recovery of chromatin surrounding early origins of DNA replication. **A**. Sequencing read depth at 114 early and 114 late replicating origins (Belsky et al. 2015) from the data described in Figure 1A (note the log2 scale). Mean sequencing coverage of early and late origins is 4163.8 and 207.7 reads respectively. *T-test* **** p ≤ 0.0001. **B and C**. Nucleosome occupancy profiles for the EdU early origin labeling experiment described in Figure 1 and for a separate experiment where following the HU-arrest, the EdU-labeled cells were allowed to proceed back into the cell cycle. Nucleosome occupancy profiles were calculated from 539 gene promoters located within 3500 bp of an early origin. **C-E**. NCOPs showing EdU labeled chromatin from arrested cells in HU and following a 2 hour release. The gene locations are the same as described in Fig 1B, C and D.

**Supplemental Figure S2.**
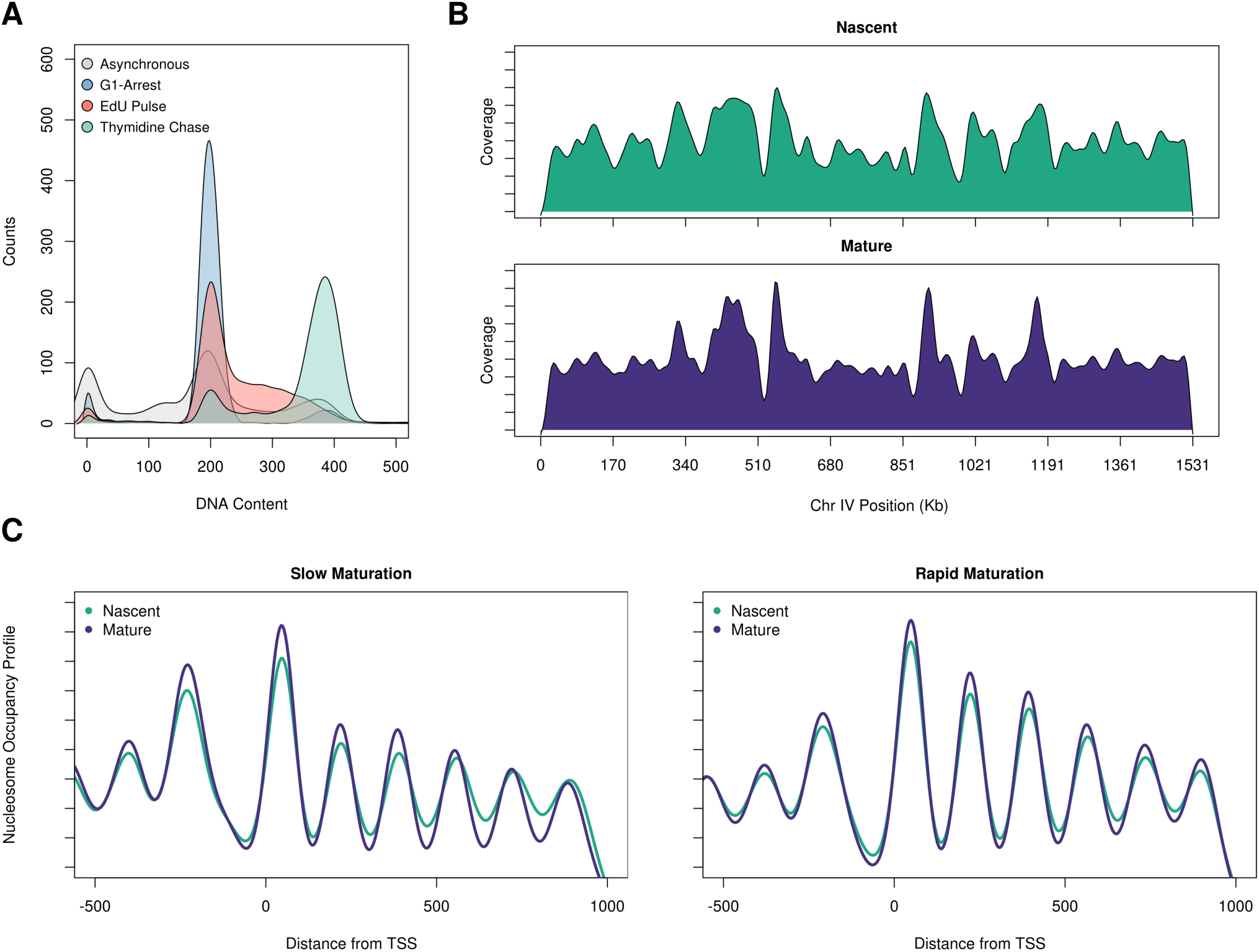
Experimental conditions to capture nascent and mature chromatin. **A**. A representative flow cytometry plot shows the cell cycle profile of yeast labeled with EdU (red track) and the state of cells following a 30 min thymidine chase (green track). **B**. Coverage plot of nascent and mature chromatin over chromosome IV shows relatively similar levels of EdU incorporation across the chromosome. **C**. Nucleosome occupancy profiles of the slow maturing (first quintile) and fast maturing (fifth quintile) genes.

**Supplemental Figure S3.**
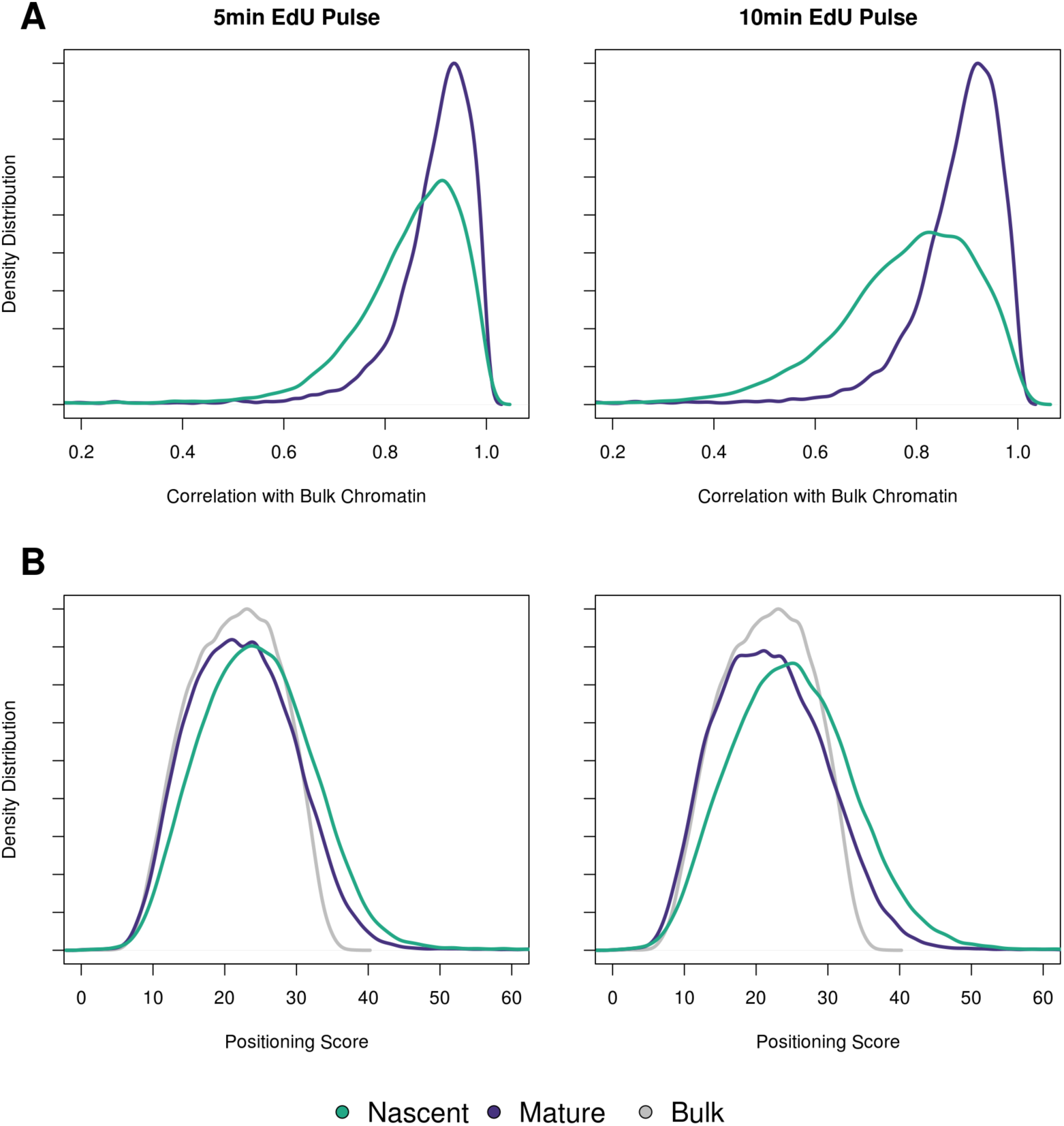
Nucleosome organization for nascent and mature chromatin from a 5 or 10 min EdU pulse. The efficacy of a 5 or 10 minute EdU pulse was assessed in replicate for each experiment. **A**. Correlation of nascent or mature chromatin with bulk chromatin from either a 5 (left) or 10 (right) min EdU pulse for gene bodies. **B**. Distribution of the nucleosome positioning scores genome-wide for nascent and mature chromatin from the 5 and 10 min EdU labeling pulse (see figure 3A).

**Supplemental Figure S4.**
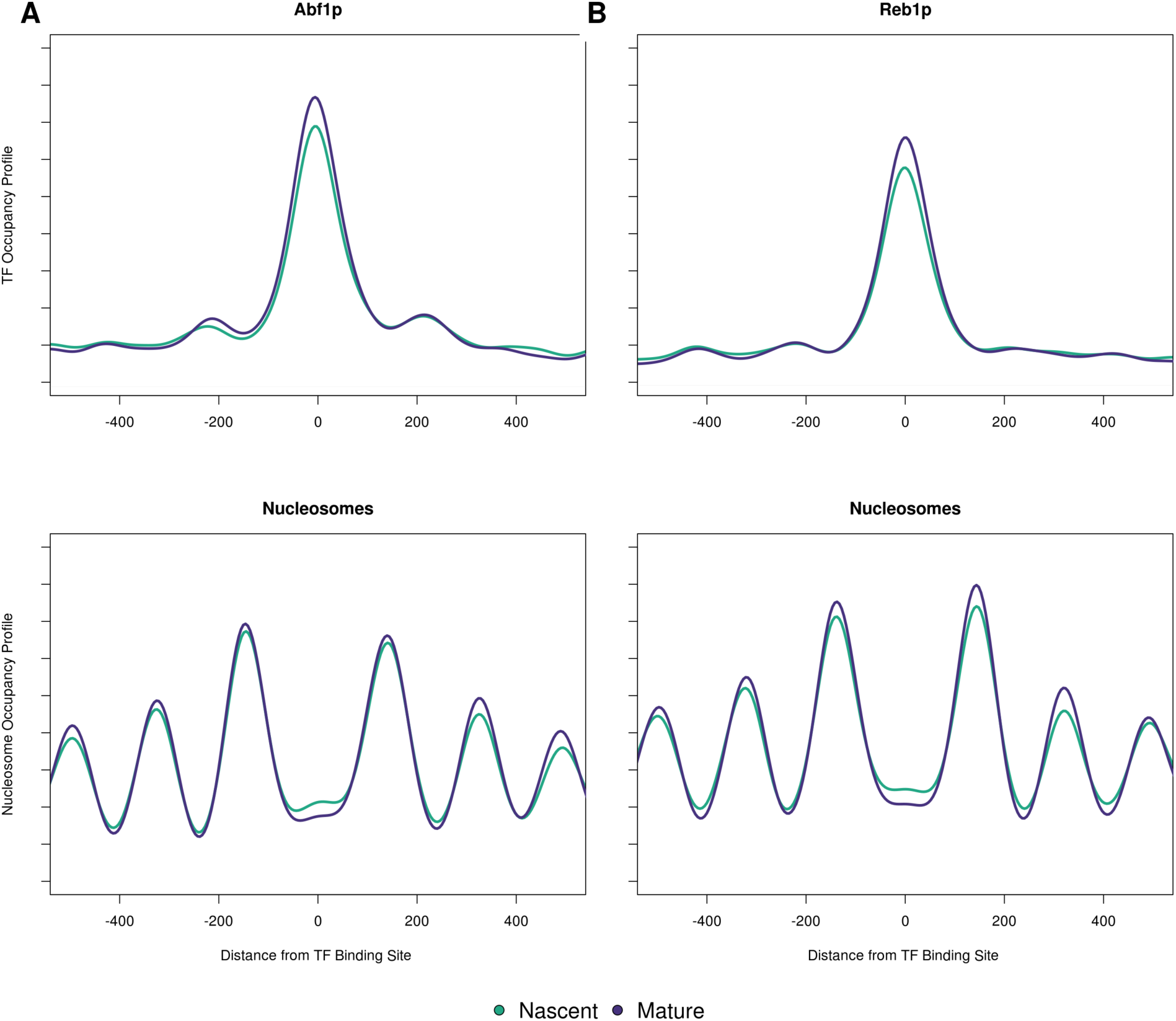
Chromatin profiles at general regulatory factor binding sites. **A.** Transcription factor occupancy for 151 Abf1p and **B**. 156 Reb1p binding sites (top two panels) and the corresponding nucleosome occupancy profiles (bottom two panels).

## References

Alabert, Constance, Jimi-Carlo Bukowski-Wills, Sung-Bau Lee, Georg Kustatscher, Kyosuke Nakamura, Flavia de Lima Alves, Patrice Menard, Jakob Mejlvang, Juri Rappsilber, and Anja Groth. 2014. “Nascent Chromatin Capture Proteomics Determines Chromatin Dynamics during DNA Replication and Identifies Unknown Fork Components.” Nature Cell Biology 16 (3): 281–93.

Alabert, Constance, and Anja Groth. 2012. “Chromatin Replication and Epigenome Maintenance.” Nature Reviews. Molecular Cell Biology 13 (3): 153–67.

Allis, C. David, and Thomas Jenuwein. 2016. “The Molecular Hallmarks of Epigenetic Control.” Nature Reviews. Genetics 17 (8): 487–500.

Annunziato, Anthony T., and Ronald L. Seale. 1982. “Maturation of Nucleosomal and Nonnucleosomal Components of Nascent Chromatin: Differential Requirements for Concurrent Protein Synthesis.” Biochemistry 21 (22): 5431–38.

Aparicio, Oscar M. 2013. “Location, Location, Location: It’s All in the Timing for Replication Origins.” Genes & Development 27 (2): 117–28.

Bai, Lu, Gilles Charvin, Eric D. Siggia, and Frederick R. Cross. 2010. “Nucleosome-Depleted Regions in Cell-Cycle-Regulated Promoters Ensure Reliable Gene Expression in Every Cell Cycle.” Developmental Cell 18 (4): 544–55.

Belsky, Jason A., Heather K. MacAlpine, Yoav Lubelsky, Alexander J. Hartemink, and David M. MacAlpine. 2015. “Genome-Wide Chromatin Footprinting Reveals Changes in Replication Origin Architecture Induced by Pre-RC Assembly.” Genes & Development 29 (2): 212–24.

Berbenetz, Nicolas M., Corey Nislow, and Grant W. Brown. 2010. “Diversity of Eukaryotic DNA Replication Origins Revealed by Genome-Wide Analysis of Chromatin Structure.” PLoS Genetics 6 (9): e1001092.

Bernstein, Bradley E., Chih Long Liu, Emily L. Humphrey, Ethan O. Perlstein, and Stuart L. Schreiber. 2004. “Global Nucleosome Occupancy in Yeast.” Genome Biology 5 (9): R62.

Boeger, Hinrich, Joachim Griesenbeck, J. Seth Strattan, and Roger D. Kornberg. 2004. “Removal of Promoter Nucleosomes by Disassembly rather than Sliding in Vivo.” Molecular Cell 14 (5): 667–73.

Breier, Adam M., Sourav Chatterji, and Nicholas R. Cozzarelli. 2004. “Prediction of Saccharomyces Cerevisiae Replication Origins.” Genome Biology 5 (4): R22.

Cayrou, Christelle, Benoit Ballester, Isabelle Peiffer, Romain Fenouil, Philippe Coulombe, Jean-Christophe Andrau, Jacques van Helden, and Marcel Méchali. 2015. “The Chromatin Environment Shapes DNA Replication Origin Organization and Defines Origin Classes.” Genome Research 25 (12): 1873–85.

Chang, L., S. S. Loranger, C. Mizzen, S. G. Ernst, C. D. Allis, and A. T. Annunziato. 1997. “Histones in Transit: Cytosolic Histone Complexes and Diacetylation of H4 during Nucleosome Assembly in Human Cells.” Biochemistry 36 (3): 469–80.

Cheloufi, Sihem, Ulrich Elling, Barbara Hopfgartner, Youngsook L. Jung, Jernej Murn, Maria Ninova, Maria Hubmann, et al. 2015. “The Histone Chaperone CAF-1 Safeguards Somatic Cell Identity.” Nature 528 (7581): 218–24.

Chen, Jiji, Andrew Miller, Ann L. Kirchmaier, and Joseph M. K. Irudayaraj. 2012. “Single-Molecule Tools Elucidate H2A.Z Nucleosome Composition.” Journal of Cell Science 125 (Pt 12): 2954–64.

Churchman, L. Stirling, and Jonathan S. Weissman. 2011. “Nascent Transcript Sequencing Visualizes Transcription at Nucleotide Resolution.” Nature 469 (7330): 368–73.

Cui, Kairong, and Keji Zhao. 2012. “Genome-Wide Approaches to Determining Nucleosome Occupancy in Metazoans Using MNase-Seq.” In Chromatin Remodeling: Methods and Protocols, edited by Randall H. Morse, 413–19. Totowa, NJ: Humana Press.

Cusick, Michael E., Paul M. Wassarman, and Melvin L. Depamphilis. 1989. “[15] Application of Nucleases to Visualizing Chromatin Organization at Replication Forks.” In Methods in Enzymology, edited by Paul M. Wassarman and Roger D. Kornberg, 170:290–316. Academic Press.

Dabin, Juliette, Anna Fortuny, and Sophie E. Polo. 2016. “Epigenome Maintenance in Response to DNA Damage.” Molecular Cell 62 (5): 712–27.

DePamphilis, M. L., and P. M. Wassarman. 1980. “Replication of Eukaryotic Chromosomes: A Close-up of the Replication Fork.” Annual Review of Biochemistry 49: 627–66.

Duronio, Robert J. 2012. “Developing S-Phase Control.” Genes & Development 26 (8): 746–50.

Eaton, Matthew L., Kyriaki Galani, Sukhyun Kang, Stephen P. Bell, and David M. MacAlpine. 2010. “Conserved Nucleosome Positioning Defines Replication Origins.” Genes & Development 24 (8): 748–53.

ENCODE Project Consortium. 2012. “An Integrated Encyclopedia of DNA Elements in the Human Genome.” Nature 489 (7414): 57–74.

Exner, Vivien, Patti Taranto, Nicole Schönrock, Wilhelm Gruissem, and Lars Hennig. 2006. “Chromatin Assembly Factor CAF-1 Is Required for Cellular Differentiation during Plant Development.” Development 133 (21): 4163–72.

Fedor, M. J., N. F. Lue, and R. D. Kornberg. 1988. “Statistical Positioning of Nucleosomes by Specific Protein-Binding to an Upstream Activating Sequence in Yeast.” Journal of Molecular Biology 204 (1): 109–27.

Fennessy, Ross T., and Tom Owen-Hughes. 2016. “Establishment of a Promoter-Based Chromatin Architecture on Recently Replicated DNA Can Accommodate Variable Inter-Nucleosome Spacing.” Nucleic Acids Research 44 (15): 7189–7203.

Ganapathi, Mythily, Michael J. Palumbo, Suraiya A. Ansari, Qiye He, Kyle Tsui, Corey Nislow, and Randall H. Morse. 2011. “Extensive Role of the General Regulatory Factors, Abf1 and Rap1, in Determining Genome-Wide Chromatin Structure in Budding Yeast.” Nucleic Acids Research 39 (6): 2032–44.

Gasser, R., T. Koller, and J. M. Sogo. 1996. “The Stability of Nucleosomes at the Replication Fork.” Journal of Molecular Biology 258 (2): 224–39.

Guillemette, Benoît, Alain R. Bataille, Nicolas Gévry, Maryse Adam, Mathieu Blanchette, François Robert, and Luc Gaudreau. 2005. “Variant Histone H2A.Z Is Globally Localized to the Promoters of Inactive Yeast Genes and Regulates Nucleosome Positioning.” PLoS Biology 3 (12): e384.

Gutiérrez, Mónica P., and David M. MacAlpine. 2016. “Chromatin Determinants of Origin Selection and Activation.” In The Initiation of DNA Replication in Eukaryotes, edited by Daniel L. Kaplan, 87–104. Cham: Springer International Publishing.

Hartley, Paul D., and Hiten D. Madhani. 2009. “Mechanisms That Specify Promoter Nucleosome Location and Identity.” Cell 137 (3): 445–58.

Hawkins, Michelle, Renata Retkute, Carolin A. Müller, Nazan Saner, Tomoyuki U. Tanaka, Alessandro P. S. de Moura, and Conrad A. Nieduszynski. 2013. “High-Resolution Replication Profiles Define the Stochastic Nature of Genome Replication Initiation and Termination.” Cell Reports 5 (4): 1132–41.

Henikoff, Jorja G., Jason A. Belsky, Kristina Krassovsky, David M. MacAlpine, and Steven Henikoff. 2011. “Epigenome Characterization at Single Base-Pair Resolution.” Proceedings of the National Academy of Sciences of the United States of America 108 (45): 18318–23.

Huang, Hongda, Caroline B. Strømme, Giulia Saredi, Martina Hödl, Anne Strandsby, Cristina González-Aguilera, Shoudeng Chen, Anja Groth, and Dinshaw J. Patel. 2015. “A Unique Binding Mode Enables MCM2 to Chaperone Histones H3-H4 at Replication Forks.” Nature Structural & Molecular Biology 22 (8): 618–26.

Ishiuchi, Takashi, Rocio Enriquez-Gasca, Eiji Mizutani, Ana Bošković, Celine Ziegler-Birling, Diego Rodriguez-Terrones, Teruhiko Wakayama, Juan M. Vaquerizas, and Maria-Elena Torres-Padilla. 2015. “Early Embryonic-like Cells Are Induced by Downregulating Replication-Dependent Chromatin Assembly.” Nature Structural & Molecular Biology 22 (9): 662–71.

Jasencakova, Zuzana, Annette N. D. Scharf, Katrine Ask, Armelle Corpet, Axel Imhof, Geneviève Almouzni, and Anja Groth. 2010. “Replication Stress Interferes with Histone Recycling and Predeposition Marking of New Histones.” Molecular Cell 37 (5): 736–43.

Jiang, Cizhong, and B. Franklin Pugh. 2009. “Nucleosome Positioning and Gene Regulation: Advances through Genomics.” Nature Reviews. Genetics 10 (3): 161–72.

Kaplan, Noam, Irene K. Moore, Yvonne Fondufe-Mittendorf, Andrea J. Gossett, Desiree Tillo, Yair Field, Emily M. LeProust, et al. 2009. “The DNA-Encoded Nucleosome Organization of a Eukaryotic Genome.” Nature 458 (7236): 362–66.

Klempnauer, K. H., E. Fanning, B. Otto, and R. Knippers. 1980. “Maturation of Newly Replicated Chromatin of Simian Virus 40 and Its Host Cell.” Journal of Molecular Biology 136 (4): 359–74.

Kliszczak, Anna E., Michael D. Rainey, Brendan Harhen, Francois M. Boisvert, and Corrado Santocanale. 2011. “DNA Mediated Chromatin Pull-down for the Study of Chromatin Replication.” Scientific Reports 1 (September): 95.

Kurat, Christoph F., Joseph T. P. Yeeles, Harshil Patel, Anne Early, and John F. X. Diffley. 2017. “Chromatin Controls DNA Replication Origin Selection, Lagging-Strand Synthesis, and Replication Fork Rates.” Molecular Cell 65 (1): 117–30.

Langmead, Ben, Cole Trapnell, Mihai Pop, and Steven L. Salzberg. 2009. “Ultrafast and Memory-Efficient Alignment of Short DNA Sequences to the Human Genome.” Genome Biology 10 (3): R25.

Lee, Cheol-Koo, Yoichiro Shibata, Bhargavi Rao, Brian D. Strahl, and Jason D. Lieb. 2004. “Evidence for Nucleosome Depletion at Active Regulatory Regions Genome-Wide.” Nature Genetics 36 (8): 900–905.

Leem, S. H., C. N. Chung, Y. Sunwoo, and H. Araki. 1998. “Meiotic Role of SWI6 in Saccharomyces Cerevisiae.” Nucleic Acids Research 26 (13): 3154–58.

Lengronne, A., P. Pasero, A. Bensimon, and E. Schwob. 2001. “Monitoring S Phase Progression Globally and Locally Using BrdU Incorporation in TK(+) Yeast Strains.” Nucleic Acids Research 29 (7): 1433–42.

Leung, Kai Him Thomas, Mohamed Abou El Hassan, and Rod Bremner. 2013. “A Rapid and Efficient Method to Purify Proteins at Replication Forks under Native Conditions.” BioTechniques 55 (4): 204–6.

Li, Bing, Samantha G. Pattenden, Daeyoup Lee, José Gutiérrez, Jie Chen, Chris Seidel, Jennifer Gerton, and Jerry L. Workman. 2005. “Preferential Occupancy of Histone Variant H2AZ at Inactive Promoters Influences Local Histone Modifications and Chromatin Remodeling.” Proceedings of the National Academy of Sciences of the United States of America 102 (51): 18385–90.

Li, Ming, Arjan Hada, Payel Sen, Lola Olufemi, Michael A. Hall, Benjamin Y. Smith, Scott Forth, et al. 2015. “Dynamic Regulation of Transcription Factors by Nucleosome Remodeling.” eLife 4 (June). https://doi.org/10.7554/eLife.06249.

Li, Qing, Hui Zhou, Hugo Wurtele, Brian Davies, Bruce Horazdovsky, Alain Verreault, and Zhiguo Zhang. 2008. “Acetylation of Histone H3 Lysine 56 Regulates Replication-Coupled Nucleosome Assembly.” Cell 134 (2): 244–55.

Lubelsky, Yoav, Takayo Sasaki, Marjorie A. Kuipers, Isabelle Lucas, Michelle M. Le Beau, Sandra Carignon, Michelle Debatisse, Joseph A. Prinz, Jonathan H. Dennis, and David M. Gilbert. 2011. “Pre-Replication Complex Proteins Assemble at Regions of Low Nucleosome Occupancy within the Chinese Hamster Dihydrofolate Reductase Initiation Zone.” Nucleic Acids Research 39 (8): 3141–55.

Luk, Ed, Ngoc-Diep Vu, Kem Patteson, Gaku Mizuguchi, Wei-Hua Wu, Anand Ranjan, Jonathon Backus, et al. 2007. “Chz1, a Nuclear Chaperone for Histone H2AZ.” Molecular Cell 25 (3): 357–68.

MacAlpine, David M., and Geneviève Almouzni. 2013. “Chromatin and DNA Replication.” Cold Spring Harbor Perspectives in Biology 5 (8): a010207.

MacIsaac, Kenzie D., Ting Wang, D. Benjamin Gordon, David K. Gifford, Gary D. Stormo, and Ernest Fraenkel. 2006. “An Improved Map of Conserved Regulatory Sites for Saccharomyces Cerevisiae.” BMC Bioinformatics 7 (March): 113.

Mantiero, Davide, Amanda Mackenzie, Anne Donaldson, and Philip Zegerman. 2011. “Limiting Replication Initiation Factors Execute the Temporal Programme of Origin Firing in Budding Yeast.” The EMBO Journal 30 (23): 4805–14.

Mathiasen, David P., and Michael Lisby. 2014. “Cell Cycle Regulation of Homologous Recombination in Saccharomyces Cerevisiae.” FEMS Microbiology Reviews 38 (2): 172–84.

Mavrich, Travis N., Ilya P. Ioshikhes, Bryan J. Venters, Cizhong Jiang, Lynn P. Tomsho, Ji Qi, Stephan C. Schuster, Istvan Albert, and B. Franklin Pugh. 2008. “A Barrier Nucleosome Model for Statistical Positioning of Nucleosomes throughout the Yeast Genome.” Genome Research 18 (7): 1073–83.

McGuffee, Sean R., Duncan J. Smith, and Iestyn Whitehouse. 2013. “Quantitative, Genome-Wide Analysis of Eukaryotic Replication Initiation and Termination.” Molecular Cell 50 (1): 123–35.

McKnight, S. L., and O. L. Miller Jr. 1977. “Electron Microscopic Analysis of Chromatin Replication in the Cellular Blastoderm Drosophila Melanogaster Embryo.” Cell 12 (3): 795–804.

Miotto, Benoit, Zhe Ji, and Kevin Struhl. 2016. “Selectivity of ORC Binding Sites and the Relation to Replication Timing, Fragile Sites, and Deletions in Cancers.” Proceedings of the National Academy of Sciences of the United States of America 113 (33): E4810–19.

Mosammaparast, Nima, Courtney S. Ewart, and Lucy F. Pemberton. 2002. “A Role for Nucleosome Assembly Protein 1 in the Nuclear Transport of Histones H2A and H2B.” The EMBO Journal 21 (23): 6527–38.

Pazin, M. J., P. Bhargava, E. P. Geiduschek, and J. T. Kadonaga. 1997. “Nucleosome Mobility and the Maintenance of Nucleosome Positioning.” Science 276 (5313): 809–12.

Probst, Aline V., Elaine Dunleavy, and Geneviève Almouzni. 2009. “Epigenetic Inheritance during the Cell Cycle.” Nature Reviews. Molecular Cell Biology 10 (3): 192–206.

Ramachandran, Srinivas, and Steven Henikoff. 2016. “Transcriptional Regulators Compete with Nucleosomes Post-Replication.” Cell 165 (3): 580–92.

Rhind, Nicholas, and David M. Gilbert. 2013. “DNA Replication Timing.” Cold Spring Harbor Perspectives in Biology 5 (8): a010132.

Richet, Nicolas, Danni Liu, Pierre Legrand, Christophe Velours, Armelle Corpet, Albane Gaubert, May Bakail, et al. 2015. “Structural Insight into How the Human Helicase Subunit MCM2 May Act as a Histone Chaperone Together with ASF1 at the Replication Fork.” Nucleic Acids Research 43 (3): 1905–17.

Santocanale, C., and J. F. Diffley. 1998. “A Mec1- and Rad53-Dependent Checkpoint Controls Late-Firing Origins of DNA Replication.” Nature 395 (6702): 615–18.

Sauer, Paul Victor, Jennifer Timm, Danni Liu, David Sitbon, Elisabetta Boeri-Erba, Christophe Velours, Norbert Mücke, et al. 2017. “Insights into the Molecular Architecture and Histone H3-H4 Deposition Mechanism of Yeast Chromatin Assembly Factor 1.” eLife 6 (March). https://doi.org/10.7554/eLife.23474.

Schlesinger, M. B., and T. Formosa. 2000. “POB3 Is Required for Both Transcription and Replication in the Yeast Saccharomyces Cerevisiae.” Genetics 155 (4): 1593–1606.

Segal, Eran, Yvonne Fondufe-Mittendorf, Lingyi Chen, Annchristine Thåström, Yair Field, Irene K. Moore, Ji-Ping Z. Wang, and Jonathan Widom. 2006. “A Genomic Code for Nucleosome Positioning.” Nature 442 (7104): 772–78.

Serra-Cardona, Albert, and Zhiguo Zhang. 2018. “Replication-Coupled Nucleosome Assembly in the Passage of Epigenetic Information and Cell Identity.” Trends in Biochemical Sciences 43 (2): 136–48.

Shibahara, K., and B. Stillman. 1999. “Replication-Dependent Marking of DNA by PCNA Facilitates CAF-1-Coupled Inheritance of Chromatin.” Cell 96 (4): 575–85.

Shirahige, K., Y. Hori, K. Shiraishi, M. Yamashita, K. Takahashi, C. Obuse, T. Tsurimoto, and H. Yoshikawa. 1998. “Regulation of DNA-Replication Origins during Cell-Cycle Progression.” Nature 395 (6702): 618–21.

Siow, Cheuk C., Sian R. Nieduszynska, Carolin A. Müller, and Conrad A. Nieduszynski. 2012. “OriDB, the DNA Replication Origin Database Updated and Extended.” Nucleic Acids Research 40 (Database issue): D682–86.

Sirbu, Bianca M., Frank B. Couch, and David Cortez. 2012. “Monitoring the Spatiotemporal Dynamics of Proteins at Replication Forks and in Assembled Chromatin Using Isolation of Proteins on Nascent DNA.” Nature Protocols 7 (3): 594–605.

Sirbu, Bianca M., Frank B. Couch, Jordan T. Feigerle, Srividya Bhaskara, Scott W. Hiebert, and David Cortez. 2011. “Analysis of Protein Dynamics at Active, Stalled, and Collapsed Replication Forks.” Genes & Development 25 (12): 1320–27.

Sirbu, Bianca M., W. Hayes McDonald, Huzefa Dungrawala, Akosua Badu-Nkansah, Gina M. Kavanaugh, Yaoyi Chen, David L. Tabb, and David Cortez. 2013. “Identification of Proteins at Active, Stalled, and Collapsed Replication Forks Using Isolation of Proteins on Nascent DNA (iPOND) Coupled with Mass Spectrometry.” The Journal of Biological Chemistry 288 (44): 31458–67.

Slattery, Matthew, Tianyin Zhou, Lin Yang, Ana Carolina Dantas Machado, Raluca Gordân, and Remo Rohs. 2014. “Absence of a Simple Code: How Transcription Factors Read the Genome.” Trends in Biochemical Sciences 39 (9): 381–99.

Smith, S., and B. Stillman. 1989. “Purification and Characterization of CAF-I, a Human Cell Factor Required for Chromatin Assembly during DNA Replication in Vitro.” Cell 58 (1): 15–25.

Sogo, J. M., H. Stahl, T. Koller, and R. Knippers. 1986. “Structure of Replicating Simian Virus 40 Minichromosomes. The Replication Fork, Core Histone Segregation and Terminal Structures.” Journal of Molecular Biology 189 (1): 189–204.

Stillman, B. 1986. “Chromatin Assembly during SV40 DNA Replication in Vitro.” Cell 45 (4): 555–65.

Tanaka, Seiji, Ryuichiro Nakato, Yuki Katou, Katsuhiko Shirahige, and Hiroyuki Araki. 2011. “Origin Association of Sld3, Sld7, and Cdc45 Proteins Is a Key Step for Determination of Origin-Firing Timing.” Current Biology: CB 21 (24): 2055–63.

Tyler, J. K., C. R. Adams, S. R. Chen, R. Kobayashi, R. T. Kamakaka, and J. T. Kadonaga. 1999. “The RCAF Complex Mediates Chromatin Assembly during DNA Replication and Repair.” Nature 402 (6761): 555–60.

Vasseur, Pauline, Saphia Tonazzini, Rahima Ziane, Alain Camasses, Oliver J. Rando, and Marta Radman-Livaja. 2016. “Dynamics of Nucleosome Positioning Maturation Following Genomic Replication.” Cell Reports 16 (10): 2651–65.

Viggiani, Christopher J., and Oscar M. Aparicio. 2006. “New Vectors for Simplified Construction of BrdU-Incorporating Strains of Saccharomyces Cerevisiae.” Yeast 23 (14-15): 1045–51.

Wang, Jie, Jiali Zhuang, Sowmya Iyer, Xinying Lin, Troy W. Whitfield, Melissa C. Greven, Brian G. Pierce, et al. 2012. “Sequence Features and Chromatin Structure around the Genomic Regions Bound by 119 Human Transcription Factors.” Genome Research 22 (9): 1798–1812.

Wasson, Todd, and Alexander J. Hartemink. 2009. “An Ensemble Model of Competitive Multi-Factor Binding of the Genome.” Genome Research 19 (11): 2101–12.

Weiner, Assaf, Tsung-Han S. Hsieh, Alon Appleboim, Hsiuyi V. Chen, Ayelet Rahat, Ido Amit, Oliver J. Rando, and Nir Friedman. 2015. “High-Resolution Chromatin Dynamics during a Yeast Stress Response.” Molecular Cell 58 (2): 371–86.

Weiner, Assaf, Amanda Hughes, Moran Yassour, Oliver J. Rando, and Nir Friedman. 2010. “High-Resolution Nucleosome Mapping Reveals Transcription-Dependent Promoter Packaging.” Genome Research 20 (1): 90–100.

Yadav, Tejas, and Iestyn Whitehouse. 2016. “Replication-Coupled Nucleosome Assembly and Positioning by ATP-Dependent Chromatin-Remodeling Enzymes.” Cell Reports 15 (4): 715–23.

Yan, Chao, Hengye Chen, and Lu Bai. 2018. “Systematic Study of Nucleosome-Displacing Factors in Budding Yeast.” Molecular Cell 71 (2): 294–305.e4.

Yarragudi, Arunadevi, Tsuyoshi Miyake, Rong Li, and Randall H. Morse. 2004. “Comparison of ABF1 and RAP1 in Chromatin Opening and Transactivator Potentiation in the Budding Yeast Saccharomyces Cerevisiae.” Molecular and Cellular Biology 24 (20): 9152–64.

Zhang, Yong, Zarmik Moqtaderi, Barbara P. Rattner, Ghia Euskirchen, Michael Snyder, James T. Kadonaga, X. Shirley Liu, and Kevin Struhl. 2009. “Intrinsic Histone-DNA Interactions Are Not the Major Determinant of Nucleosome Positions in Vivo.” Nature Structural & Molecular Biology 16 (8): 847–52.

